# The SPATA5-SPATA5L1-CINP-C1ORF109 complex is essential for cytoplasmic pre-60S ribosome maturation

**DOI:** 10.1101/2025.11.07.687187

**Authors:** Jonas Walter, Melanie Weisser, Magdalena Masternak, Thomas Wild, Ricardo Garcia-Martin, Tim Liebner, Elisabeth Bragado-Nilsson, Takeo Narita, Niki Zavogianni, Guillermo Montoya, Chunaram Choudhary

**Affiliations:** Department of Proteomics, The Novo Nordisk Foundation Center for Protein Research, Faculty of Health and Medical Sciences, University of Copenhagen, Blegdamsvej 3B, 2200 Copenhagen, Denmark; Structural Molecular Biology Group, Novo Nordisk Foundation Centre for Protein Research, Department of Cellular and Molecular Medicine, Faculty of Health and Medical Sciences, University of Copenhagen, Blegdamsvej 3B, Copenhagen, 2200, Denmark

**Keywords:** Ribosome, pre-60S maturation, SPATA5, SPATA5L1, CINP, C1ORF109, 55LCC

## Abstract

The ribosome is a universally conserved and essential protein complex, but its biogenesis in mammals is more complex than in single-celled eukaryotes. To explore this added complexity, we conducted a protein–protein interaction screen in human cells. This led to the identification of the eumetazoan-specific SPATA5–SPATA5L1–CINP–C1ORF109 (55LCC) complex as a key regulator of ribosome biogenesis. Structural analyses using cryo-EM and X-ray crystallography defined the architecture of 55LCC. Functional studies following acute depletion revealed that each component is essential for pre-60S maturation. Swapping endogenous 55LCC components with mutant versions pinpointed critical functional interactions and showed that SPATA5’s ATPase activity is more important than SPATA5L1’s. Our findings support that SPATA5 evolved from the solitary yeast ATPase Drg1 into the multiprotein 55LCC complex in metazoans. This work provides insights into the complexity of ribosome biogenesis and lays the foundation for deeper exploration of 55LCC’s role in pre-60S maturation.

## Introduction

Ribosomes are essential, universally conserved molecular machines that translate genetic information into functional proteins. They are highly abundant, with an estimated ∼10⁷ ribosomes per human cell, and ribosomal proteins account for ∼4-6% of the total cellular protein mass^1^, highlighting its fundamental importance. Ribosome production is estimated to consume ∼30% of cellular energy budget^2^.

The mature mammalian 80S ribosome consists of ∼80 proteins and four ribosomal RNAs (18S, 5S, 5.8S, and 28S). Ribosome biogenesis is a multi-step process that begins in the nucleolus, where ribosomal RNA (rRNA) genes are transcribed by RNA polymerase I, producing precursor rRNAs (pre-rRNAs) that undergo extensive processing and modification^3^. Simultaneously, ribosomal proteins (RPs) and ribosome biogenesis factors (RBFs) are synthesized in the cytoplasm and imported into the nucleolus, where they assemble into the 90S pre-ribosome. This complex subsequently splits into pre-40S and pre-60S particles, which follow independent maturation pathways. The final steps of ribosome biogenesis occur in the cytoplasm, culminating in the joining of 40S and 60S and the formation of functional 80S ribosomes^4^.

Budding yeast has served as the key model system for studying eukaryotic ribosome biogenesis, leading to the identification of nearly 200 trans-acting RBFs^5^. However, increasing evidence suggests that mammalian ribosome biogenesis is more complex, involving additional auxiliary factors^6^. Some of these higher metazoan-specific RBFs are linked to ribosomopathies, diseases arising from defects in ribosome assembly or function^7^.

To explore the intricacy of mammalian ribosome biogenesis, we performed affinity enrichment and mass spectrometry (AP-MS) analysis on key endogenously expressed RBFs in human cells. In addition to detecting many known interactors, our screen identified putatively new interactors, amongst AAA+ ATPases SPATA5 and SPATA5L1 as interactors of the GTPase GTPBP4 (Nog1). Reverse PPI analyses of SPATA5 and SPATA5L1 confirmed their link to GTPBP4 and identified CINP and C1ORF109 as additional interactors. Biochemical and functional analyses confirmed that SPATA5, SPATA5L1, CINP, and C1ORF109 form a tightly interacting complex. During the course of this work, three groups identified this complex and named it 55LCC^8–10^. Our findings support the reported structure of 55LCC^8,10^ and its role in ribosome biogenesis^9^ while offering complementary new insights into its function in pre-60S maturation.

## Results

### Mapping human ribosome-interacting proteins

For our AP-MS screen, we selected six RPs and 12 known RBFs involved in regulating different steps of ribosome maturation. For detecting 90S interactions we used CIRH1A/UTP4, UTP6, UTP11, UTP18 and NOP56^11^. For detecting pre-40S interactions, we used NOB1 and TSR1^11^. For identifying pre-60S interactors we used PES1, GTPBP4 (Nog1) and PA2G4/EBP1^11^. For detecting 80S/60S/40S interactions we used uL10/RPLP0, uL1/RPL10A, eL28/RPL28, uS3/RPS3, eS1/RPS3A and uS5/RPS2^12^.

To study interactions under native expression conditions, 18 proteins were endogenously tagged in the human HCT116 cell line(**Figure 1A, Supplementary Figure S1A, Supplementary Table 1**). All these proteins are common essential for human cells, their homozygous tagging suggest that they retained their function. Expression and subcellular localization of tagged bait proteins were confirmed using immunofluorescence. Different tagged bait proteins showed distinct subcellular localization, consistent with their reported roles in regulating different steps of ribosome biogenesis or available imaging data (**Figure 1B**, **Supplementary Figure S1B**).

**Figure 1.**
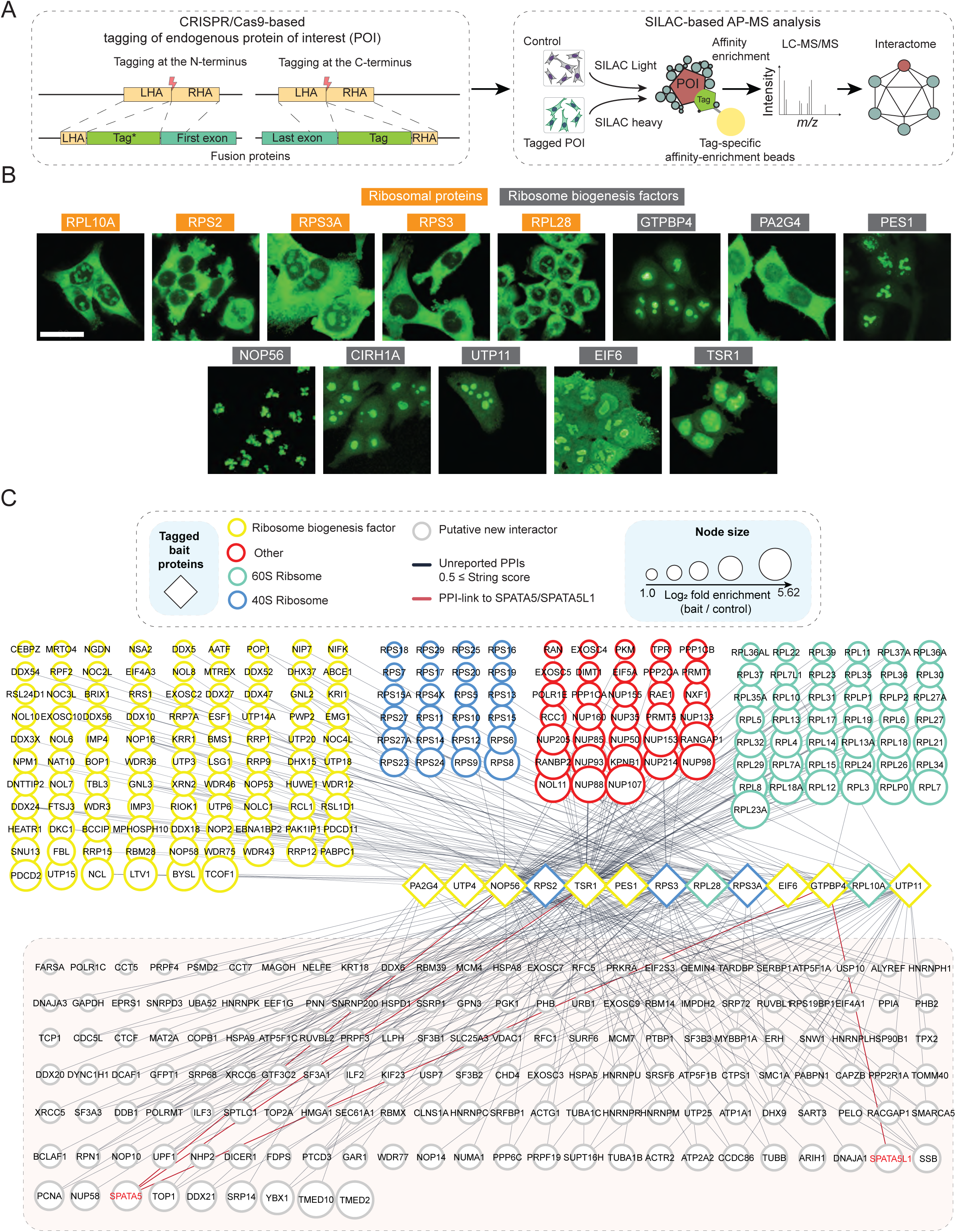
Identification of ribosome biogenesis-associated factors. (**A**) Strategy for CRISPR/Cas9-based gene editing to generate endogenously tagged protein of interest (POI, baits) (left panel) in human HCT116 cells. Schematic for affinity-purification mass spectrometry (AP-MS) analysis of affinity-tagged factors (right panel). (**B**) Micrographs showing subcellular distribution of 13 bait proteins used for AP-MS analyses (also see **Supplementary Figure S1**). (**C**) Interaction network of proteins enriched with the specified bait proteins, visualized using Cytoscape. AP-MS analyses (*n*≥3 per bait) were performed using the indicated bait proteins, including ribosomal proteins and ribosome biogenesis factors. The network encodes multiple data dimensions: nodes represent individual interactors, and node size reflects the relative enrichment strength (maximum value per screen). Known ribosomal interactors were categorized and color-coded as indicated. Edges represent protein–protein interactions (PPIs) among significantly enriched proteins with a STRING interaction score <0.5. Edges and nodes corresponding to our follow-up candidates are highlighted in red. Unconnected nodes correspond to proteins with STRING scores ≥0.5 but not listed among manually curated known ribosome biogenesis factors (**Supplementary Table 4**). All identified interactors are listed in **Supplementary Table 8,** and the processed data for network generation are in **Supplementary Table 9**.

Using this set of baits, we performed affinity-enrichment and mass spectrometry (AP-MS) to identify their interactors. Using SILAC-based quantification, enrichment of bait-associated proteins was compared with mock pull-down from parental cells (**Supplementary Figure S2**), similar to previous work in yeast^13^. While some baits (e.g., GTPBP4, PA2G4, EIF6, PES1) interacted with many known RBFs and RPs, others (i.e., UTP6, UTP18, NOB1, MDN1, RPLP0) did not show a notable enrichment of RPs and RBFs, and due to this, the interactome data were not processed (**Supplementary Figure S2, Supplementary Table 2**). To identify new candidate interactors involved in ribosome biogenesis, we posited that such new interactors must be essential for cell viability, as ribosome biogenesis is essential for cell proliferation. We defined essential proteins/genes as those required for proliferation for >=50% (0.5 score) of human cell lines listed in the DepMap database^14^ (**Supplementary Table 3**). Interactions among enriched proteins were identified using the STRING database^15^ (**Figure 1C**). Those essential interactors with STRING interaction scores <0.5, while not being listed among known RBFs (**Supplementary Table 4**), were considered putative new interactors. These were ranked based on their enrichment level in our AP-MS analyses (**Figure 1C, grey circles**).

### SPATA5, SPATA5L1, CINP, and C1ORF109 form a multi-protein complex

Among the top enriched candidates not previously linked to ribosome biogenesis, SPATA5 and SPATA5L1 stood out. Both were significantly and strongly enriched in GTPBP4 interactomes, are among essential human proteins in the DepMap database, and possess predicted ATPase domains, making them interesting candidates for further investigation. To confirm these interactions, endogenous SPATA5 and SPATA5L1 were fused with GFP using CRISPR-based genome editing in HCT116 cells. These reciprocal AP-MS analyses of SPATA5 and SPATA5L1 confirmed the connection with GTPBP4 (**Figure 2A**) and, interestingly, strong mutual enrichment.

**Figure 2.**
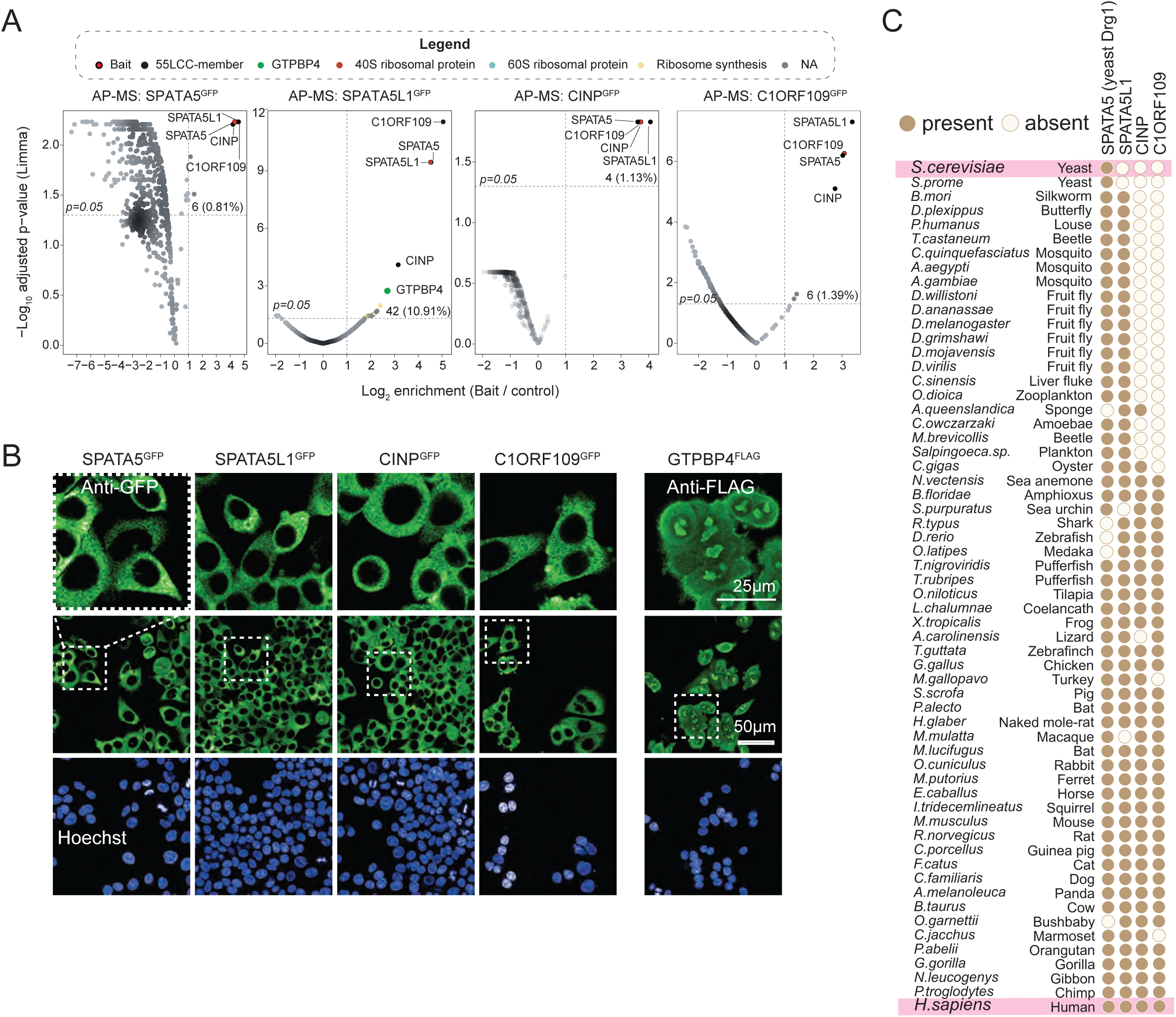
Identification of the ribosome-interacting 55LCC complex. (**A**) AP-MS analysis of endogenously expressed SPATA5, SPATA5L1, CINP, and C1ORF109. Endogenous genes expressing the indicated proteins were fused with GFP using CRISPR-based genome editing. Volcano plots show proteins enriched in the indicated AP-MS analysis. Statistical significance determined using LIMMA^61^. Each plot indicates the bait protein and the total number (*n*) of significantly enriched proteins (top). A horizontal line marks the p = 0.05 significance threshold. (**B**) Micrographs showing the subcellular localization of the endogenously expressed GFP-fused SPATA5, SPATA5L1, CINP, and C1ORF109 used for AP-MS. For comparison subcellular localization of endogenously expressed FLAG-tagged GTPBP4, a pre-60S associated biogenesis factor and an interactor of the 55LCC, is shown. (**C**) Evolutionary conservation of genes 55LCC components across selected species. Orthologs of human SPATA5, SPATA5L1, CINP, and C1ORF109 were identified in the indicated species using a reciprocal BLAST search.

The yeast ortholog of SPATA5, Drg1, is known to be required for the maturation of pre-60S in the cytoplasm^16,17^. The ATPase activity of Drg1 is required for the release of Rlp24 (RSL24D1), eIF6 (EIF6), and Nog1 (GTPBP4) from the pre-60S. In our analyses, SPATA5/SPATA5L1 interactors also included EIF6 and GTPBP4, supporting a conserved role in pre-60S maturation.

Notably, both SPATA5 and SPATA5L1 showed strong interactions with CINP (cyclin-dependent kinase 2 interacting protein) and C1ORF109, a protein of unknown function. Although both CINP and C1ORF109 are listed as essential genes in the DepMap database, neither had previously been associated with ribosome biogenesis. To validate these interactions, we endogenously tagged CINP and C1ORF109 with GFP using CRISPR-based genome editing. Reciprocal AP-MS analyses of CINP and C1ORF109 showed that these four proteins (SPATA5, SPATA5L1, CINP, and C1ORF109) interact with each other (**Figure 2A**), suggesting they form a complex, hereafter referred to as 55LCC^8^.

Subcellular localization analyses show that all four members of 55LCC are almost exclusively present in the cytoplasm (**Figure 2B**). In contrast, interacting GTPBP4 prominently localizes across the nucleolus, nucleoplasm, and cytoplasm. This is consistent with the data that GTPBP4 binds to pre-60S in the nucleolus and remains attached to it until the final steps of pre-60S maturation in the cytoplasm^18^. The cytoplasmic localization of 55LCC is consistent with the localization of Drg1 in yeast^16,19^, suggesting that 55LCC likely associates with the pre-60S in the cytoplasm.

Next, we examined the presence of 55LCC constituents in evolutionarily diverse animals. Sequence homology analysis of all four proteins revealed that SPATA5 is conserved across all eukaryotes, SPATA5L1 evolved in early metazoans, and CINP and C1ORF109 evolved subsequently (**Figure 2C**).

### Acute depletion of 55LCC constituents arrests cell proliferation

To study the functional roles of the 55LCC constituents, we generated cell lines to acutely deplete them. Using CRISPR-based editing, each of the genes encoding 55LCC constituents was fused with the FKBP12^F36V^ degron tag (hereafter dTAG)^20^ (**Figure 3A**). To induce conditional depletion, the edited cell lines were treated with dTAG^V^-1, which triggers proteasome-mediated degradation of dTAG-fused proteins. Previous work has demonstrated that dTAG^V^-1 does not cause depletion or functional effects in cells lacking the FKBP12F36V degron, thus confirming specificity and absence of off-target effects^21^.

**Figure 3.**
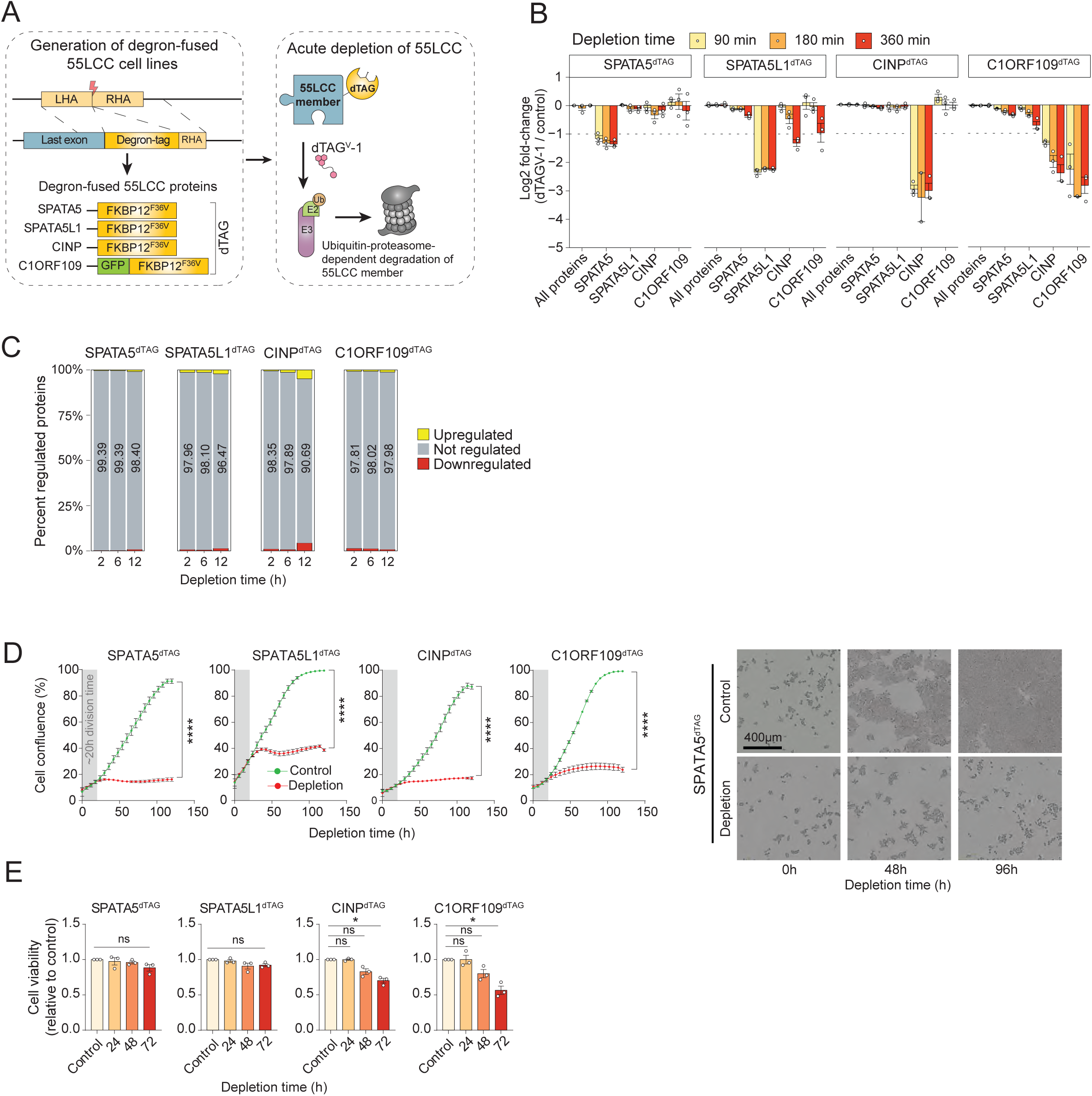
Acute depletion of 55LCC constituents causes cell proliferation arrest. (**A**) Schematic of CRISPR-based generation of SPATA5, SPATA5L1, CINP, and C1ORF109 depletion cell lines. Using CRISPR, endogenous genes were fused with the FKBP12^F36C degron tag (dTAG). dTAG^V^-1 treatment triggers acute depletion of dTAG-fused proteins through the ubiquitin-proteasome system. (**B**) Validation of degron-fused 55LCC constituents. Degron-fused 55LCC expressing cells (from A) were treated with 50nM dTAG^V^-1 (standard concentration in all experiments) for the indicated time, and changes in protein expression quantified using mass spectrometry (*n*=3). Shown are fold-changes in 55LCC expression after the depletion of specified proteins. ‘All proteins’ indicate median fold changes of all quantified proteins. (**C**) Quantification of global protein changes after 55LCC depletion. The specified 55LCC complex members were acutely depleted for the indicated time, and the resulting changes in global protein levels quantified by mass spectrometry (*n*=3). The fraction of regulated proteins is shown. (**D**) 55LCC depletion arrests cell proliferation. Cells expressing dTAG-fused 55LCC complex members were subjected to the indicated treatments, and cell proliferation analyzed by monitoring cell confluency by live-cell imaging (by using Incucyte) (*n*=3). Shown are the changes (Mean ± SEM) in cell confluency over 150 hours (left panels). Representative micrograph images from SPATA5^dTAG^-depleted cells are shown (right panels). Statistical significance (2-way-ANOVA, Šidák-corrected): * p<0.05, ** p<0.01, *** p<0.001, **** p<0.0001. (**E**) Quantification of viable cells at indicated time points was performed (*n*=3) (Mean ± SEM, Student T-test).

To investigate the interdependency of 55LCC constituents for their stability, we individually depleted each component and monitored protein levels of all complex members, as well as global proteomic changes. Protein abundance changes were quantified at 1.5, 3, and 6 hours following dTAG^V^-1 treatment using mass spectrometry (MS) (**Figure 3B**). dTAG^V^-1 treatment strongly decreased the abundance of the targeted 55LCC member within 1.5 hours (**Figure 3B**). SPATA5L1 depletion reduced the abundance of other complex members, with a notable decrease in CINP and C1ORF109 levels. While C1ORF109 depletion strongly reduced levels of CINP, the contrary was not true. CINP was depleted efficiently, but C1ORF109 expression remained unchanged. SPATA5 depletion also did not affect levels of other interactors, although we note that SPATA5 depletion was somewhat less pronounced compared to other depletions.

The observed codependence of 55LCC proteins for their cellular stability is consistent with our *in vitro* experiments (**Supplementary Figure S3A–S3C**). In these experiments, CINP, C1ORF109, or both were co-overexpressed in *E. coli*, while SPATA5, SPATA5L1, or combinations of CINP/C1ORF109/SPATA5/SPATA5L1 were transiently co-overexpressed in human cells as outlined in **Supplementary Figure S3A** and shown by western blot analysis (**Supplementary Figure S3B**). The experiments revealed that only SPATA5 and CINP could be stably overexpressed and purified individually at high yields, whereas SPATA5L1 and C1ORF109 degraded rapidly unless co-expressed with SPATA5 and/or CINP (**Supplementary Figure S3C**). Notably, we could reconstitute stable wild-type SPATA5–SPATA5L1–CINP–C1ORF109 heterooligomers *in vitro* by affinity purification from a single tagged subunit – SPATA5 (**Supplementary Figure S3C**) and CINP, C1ORF109, or SPATA5L1 (data not shown) – in agreement with our preceding *in vivo* pull-down and MS-interactome results. In addition, we purified C1ORF109 and CINP as a stable heterodimer, with C1ORF109 stability matching that of CINP (**Figure 5B**).

To further understand the impact of 55LCC constituent depletion on global protein expression, we performed extended time-course analysis, and changes in protein expression were quantified at 2, 6, and 12 hours after inducing the depletion of each of the 55LCC constituents. Up until 12 hours, no major changes in proteome expression were observed, except for CINP, where protein expression changes became evident after 12 hours (**Figure 3C**).

Imaging-based automated growth analysis revealed that long-term depletion of each 55LCC complex member led to arrest in cell proliferation (**Figure 3D, Supplementary Figure S3D**). However, the reduction in cell proliferation only starts to become evident after one day, which roughly corresponds to the doubling time of HCT116 cells. Analysis of cell viability and morphology after 55LCC depletion indicates that cells remain viable and attached to culture dishes for at least 24 hours after inducing the depletion, but at later points, cells start to die (**Figure 3E**, **Supplementary Figure S3D**). Cell death is more pronounced in CINP^dTAG^ and C1ORF109^dTAG^ depleted cells, maybe due to the more complete depletion of these proteins.

Together, these results confirmed selective depletion of dTAG-fused 55LCC constituents and individual critical importance for cell proliferation and viability.

### Acute 55LCC depletion impairs ribosome assembly and protein synthesis

To directly investigate 55LCĆs involvement in ribosome biogenesis, we analyzed interactions of 60S ribosomal subunits, with and without depleting 55LCC constituents (**Supplementary Figure S4A**). We chose RPL10A and RPL28 for these analyses as both are present in pre-60S as well as in mature 80S ribosomes. Also, it has been demonstrated that the C-terminal GFP tagging of RPL28^9^ and N-terminal tagging of RPL10A^22^ do not interfere with their incorporation into the ribosome. In the SPATA5^dTAG^ and CINP^dTAG^ cell lines, RPL10A and RPL28 were fused with GFP endogenously.

Using these cell systems, we analyzed ^GFP^RPL10A and RPL28^GFP^ interactions, with and without depleting SPATA5^dTAG^ or CINP^dTAG^ (**Supplementary Figure S4B**). Notably, in both ^GFP^RPL10A and RPL28^GFP^ interactomes, GTPBP4, RSL24D1, EIF6, and MRTO4 are enriched significantly more strongly after depletion of either SPATA5^dTAG^ or CINP^dTAG^ (**Figure 4A**), suggesting an accumulation of the pre-60S bound to these proteins.

**Figure 4.**
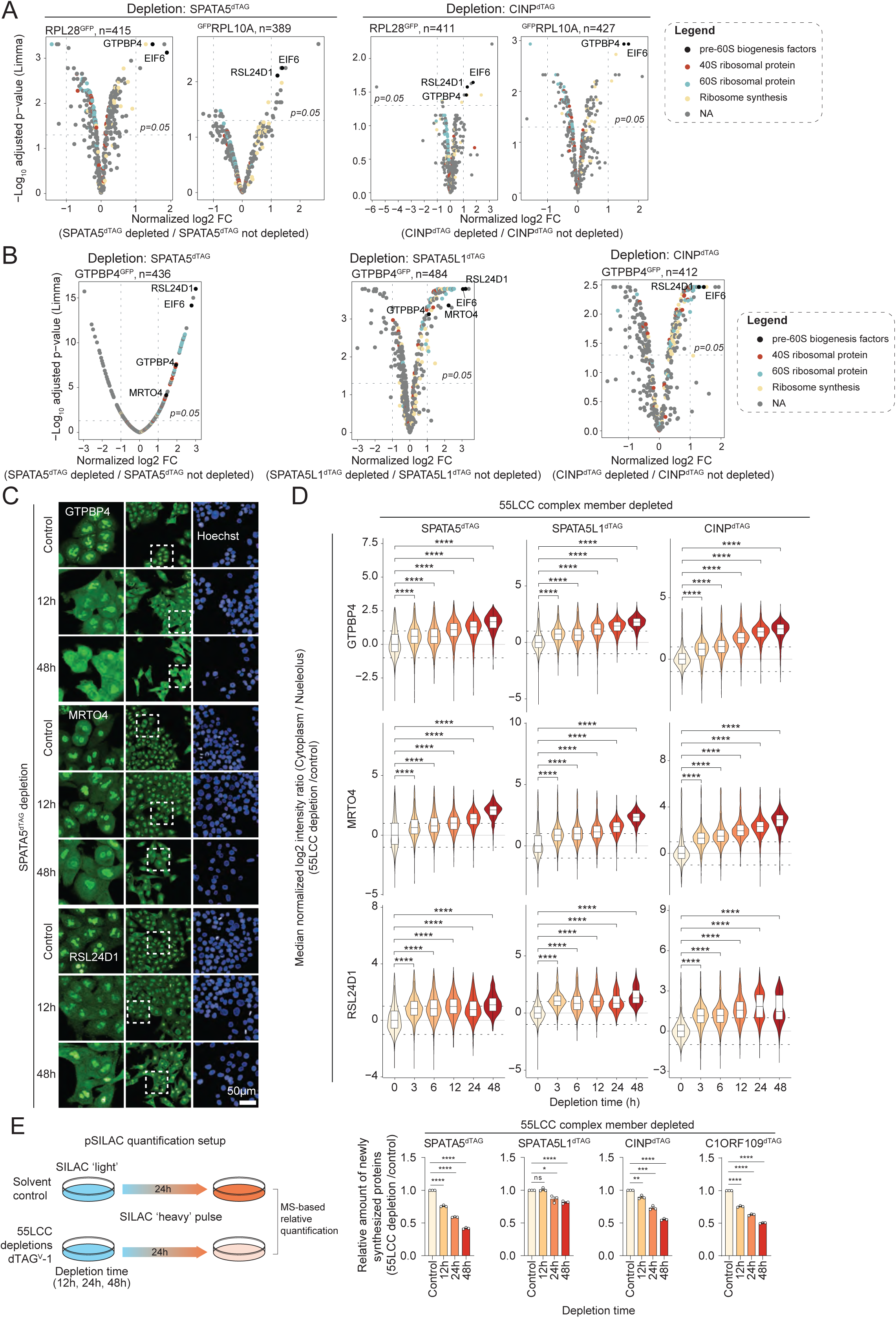
Acute depletion of 55LCC impairs pre-60S maturation and reduces protein synthesis. (**A**) Analysis of ribosome-associated interactors after the depletion of 55LCC. Endogenous RPL10A and RPL28 were fused with GFP in SPATA5^dTAG^ and CINP^dTAG^ expressing cells. ^GFP^RPL10A and RPL28^GFP^ interactions were analyzed by AP-MS, with and without acute depletion of SPATA5^dTAG^ and CINP^dTAG^. Enrichment of identified interactors is compared between control and SPATA5^dTAG^ and CINP^dTAG^ depleted cells. Selected pre-60S-associated ribosome biogenesis factors enriched after SPATA5^dTAG^ and CINP^dTAG^ depletion are shown. (*n*=3) (**B**) Endogenous GTPBP4 was fused with GFP in SPATA5^dTAG^, SPATA5L1^dTAG^ and CINP^dTAG^ expressing cells, and GTPBP4^GFP^ interacting proteins were identified with or without depletion of the specified 55LCC constituents, as described in A (*n*=3). (**C**) Representative micrographs showing the subcellular localization of GTPBP4, MRTO4 and RSL24D1 in the indicated conditions. GTPBP4, MRTO4 and RSL24D1 were stained with or without depletion of SPATA5^dTAG^ for the specified time. (**D**) Quantification of fluorescence intensities of the indicated pre-60S biogenesis factors in specified subcellular compartments at the indicated timepoints after the depletion of SPATA5^dTAG^, SPATA5L1^dTAG^ or CINP^dTAG^ (*n*=3). Violin plots display the distribution of log₂-transformed intensity ratios between cytoplasm and nucleus across treatment time points. Boxplots are overlaid within each plot and indicate the median (horizontal bar) and the interquartile range (IQR; box hinges at the 25th and 75th percentiles). Whiskers extend to 1.5× the IQR; outliers beyond this range are not shown. Statistical comparisons to the untreated control (0 h) were performed using a two-sided Wilcoxon test with Benjamini– Hochberg correction for multiple testing **** p<0.0001. Dotted lines at ±1 log2 fold change. (**E**) Setup of pulsed-SILAC-based quantification of newly synthesized proteins after acute depletion of 55LCC (left panel). Relative protein synthesis rate (right panel) after the depletion of the indicated dTAG-fused 55LCC constituents, for the specified time (*n*=3) (Mean ± SEM) (Student T-test; * p<0.05, ** p<0.01, *** p<0.001, **** p<0.0001, ns: not significant).

**Figure 5.**
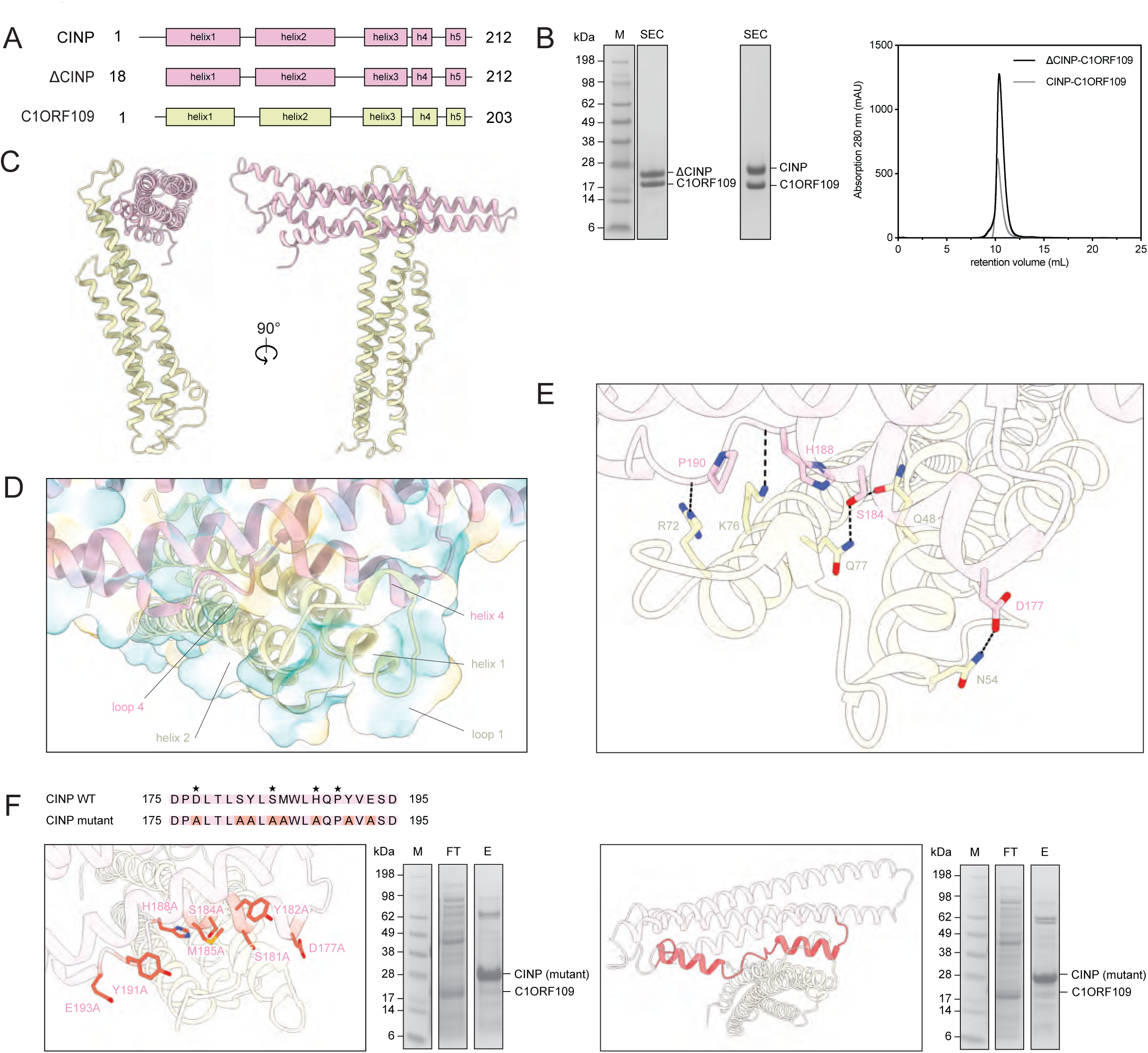
Crystal structure of the ΔCINP-C1ORF109 dimer. (**A**) Secondary structure schematic of proteins CINP, ΔCINP, and C1ORF109. (**B**) **Left panel:** Coomassie-stained SDS-PAGE analysis of purified selenomethionine-incorporated ΔCINP-C1ORF109 and native full-length CINP-C1ORF109 complexes. **Right panel:** Size-exclusion chromatography profiles of the corresponding purifications, using a Superdex 75 10/300 column. (**C**) Crystal structure of the ΔCINP-C1ORF109 complex, shown in cartoon representation. ΔCINP and C1ORF109 are colored in pink and lime, respectively. (**D**) Hydrophobic surface representation of the interacting helices and loops within the ΔCINP-C1ORF109 complex. The molecular surface is colored according to its hydrophilic and hydrophobic properties, ranging from teal (hydrophilic) to gold (hydrophobic). (**E**) Close-up view of the interaction site between ΔCINP and C1ORF109, located between helix 4 and loop 4 of ΔCINP and helices 1 and 2, as well as loop 1 of C1ORF109. Interacting residues are labeled and shown in stick representation. Hydrogen bond interactions are depicted as black dashed lines. Atoms are colored according to type: carbon in pink (ΔCINP) and lime (C1ORF109), nitrogen in blue and oxygen in red. (**F**) Mutagenesis study of CINP-C1ORF109 complex (left panel: point mutant, right panel: truncation mutant). Sequence alignment of the interacting region, highlighting mutated or truncated residues (in red) designed to disrupt the binding interface. Residues involved in interactions between CINP and C1ORF109 are marked with a black star (top panel). Close-up view on mutated residues (shown in stick representation, colored according to atom type: carbon atoms in tomato, nitrogen atoms in blue and oxygen atoms in red). Coomassie-stained SDS-PAGE analysis of the purified mutant complex. M – marker, FT – flow-through, E – elution.

To further strengthen this result, we used SPATA5^dTAG^, SPATA5L1^dTAG^, and CINP^dTAG^ expressing cells, and in these backgrounds, fused GTPBP4 with GFP endogenously. In each of the cell lines, GTPBP4^GFP^ interactomes were analyzed with and without depleting SPATA5^dTAG^, SPATA5L1^dTAG^, or CINP^dTAG^. Reassuringly, RSL24D1, EIF6, and MRTO4 show increased enrichment in GTPBP4 pull-downs after SPATA5^dTAG^, SPATA5L1^dTAG^, or CINP^dTAG^ depletion (**Figure 4B**).

Together, these unbiased depletion interactomes after 55LCC constituent depletion show a common signature and highlight that 55LCC is required for the release of pre-60S-associated factors during the final steps of 60S maturation in the cytoplasm.

To confirm this observation and to learn about the dynamics, SPATA5^dTAG^, SPATA5L1^dTAG^, and CINP^dTAG^ were depleted by dTAG^V^-1 treatment, and subcellular localization of GTPBP4, RSL24D1 and MRTO4 was analyzed 3, 6, 12, 24, and 48 hours after inducing the depletion. The results show that depletion of SPATA5^dTAG^, SPATA5L1^dTAG^, and CINP^dTAG^ leads to increased accumulation of GTPBP4, RSL24D1, and MRTO4 in the cytoplasm (**Figure 4C-D**, **Supplementary Figure S4C-D**). The results imply that GTPBP4, RSL24D1, and MRTO4 can correctly assemble onto the pre-60S subunit in the nucleus and be exported to the cytoplasm; however, they fail to be released from the pre-60S in the cytoplasm. We also examined the RPL10A localization after depleting SPATA5^dTAG^ or CINP^dTAG^. Interestingly, depletion of SPATA5^dTAG^ and CINP^dTAG^ caused an increase in RPL10A in the nucleolus and a reduction in the cytoplasm (**Supplementary Figure S5A-B**). The reason for this increased nucleolar localization remains to be investigated. One possibility is that nucleolar recycling of pre-60S-bound factors that are released by 55LCC in the cytoplasm is important for producing export-competent pre-60S.

To evaluate how depletion of 55LCC proteins affects the distribution of ribosomal subunits, we compared polysome profiles between cells expressing endogenous 55LCC proteins and cells in which 55LCC proteins were depleted. Depletion resulted in a decrease in the 80S ribosomes peak (**Supplementary Figure S5C**). Immunoblot analysis of the 40S, 60S, and 80S fractions using an antibody against GTPBP4 revealed that while GTPBP4 was absent from the 60S fraction under endogenous conditions, depletion of 55LCC constituents led to its presence in the 60S fraction. These results corroborate previous findings^9^, demonstrating that depletion of C1ORF109 and SPATA5L1 subunits disrupts ribosome biogenesis. Our analysis extends these data by examining the full complement of 55LCC constituents, including SPATA5 and CINP, and by directly monitoring GTPBP4 localization across ribosomal fractions. Notably, while the previous study observed polysome profile abnormalities and half-mer formation indicative of 60S subunit deficiency, our immunoblot analysis provides direct evidence for the accumulation of GTPBP4-loaded pre-60S particles upon 55LCC depletion. The aberrant retention of GTPBP4 in the 60S fraction (absent in wild-type cell line) supports a model in which 55LCC proteins facilitate the release of GTPBP4 from maturing pre-60S subunits, thereby preventing the accumulation of stalled ribosomal intermediates. This finding is consistent with the accumulation of GTPBP4-bound pre-60S particles upon 55LCC depletion observed in our AP-MS experiments.

To directly and globally assess how impaired ribosome biogenesis affects protein synthesis, we employed pulse SILAC^23^ to compare synthesis rates in cells with or without 55LCC depletion. Cells were pretreated with dTAG^V^-1 for 12, 24, or 48 hours (**Figure 4E**), followed by a pulse with stable isotope-labeled amino acids. Relative protein synthesis was quantified by measuring the incorporation of heavy-labeled amino acids and comparing their abundance between control and 55LCC-depleted conditions. Depletion of 55LCC components led to a marked reduction in nascent protein synthesis.

Together, AP-MS analyses with multiple baits, immunostaining of pre-60S–associated factors, and polysome profiling reveal that rapid 55LCC depletion causes accumulation of pre-60S particles associated with GTPBP4, RSL24D1, EIF6, and MRTO4. This aberrant retention of assembly factors disrupts pre-60S maturation within hours and ultimately impairs 80S replenishment and global protein synthesis.

### Crystal structure of the ΔCINP-C1ORF109 complex

We determined the structure of the ΔCINP-C1ORF109 complex (**Figure 5A**). The N-terminal tail of CINP was truncated, as a limited proteolysis assay combined with MS suggested that it was disordered and may hinder crystallization (**Supplementary Figure S6A**). The heterodimeric ΔCINP-C1ORF109 complex was crystallized, and its structure was determined to 3.69 Å using single-wavelength anomalous dispersion (SAD) (**Figure 5C**, **Supplementary Figure S6B**, **Supplementary Table 5**).

ΔCINP-C1ORF109 crystallized as a dimer of dimers, with four molecules in the asymmetric unit (chains A and C corresponding to C1ORF109, and chains B and D corresponding to ΔCINP) (**Supplementary Figure S6C**). However, the crystal asymmetric unit did not represent the active biological unit, which is formed between chain C of one asymmetric unit with chain D from a symmetry-related molecule (**Supplementary Figure S6D**). The refined structure revealed that both proteins consist of five helices, forming a coiled-coil structure (**Figure 5C**, **Supplementary Figure S6E**). Analysis of the interacting helices and loops indicated primarily hydrophobic interactions within the heterodimer (**Figure 5D**). These interactions are further stabilized by hydrogen bonds between specific residues (D177 and N54, S184 and Q48/Q77, H188 and K76, and P190 and R72 of CINP and C1ORF109) (**Figure 5E**). To assess the importance of the interaction site between CINP and C1ORF109 in the context of the 55LCC complex stability and cell viability, disruption mutants were designed (**Figure 5F**). Specifically, D177, S181, Y182, S184, M185, H188, Y191 and E183 in CINP were mutated to alanine. In addition, a truncation mutant lacking the entire G173-L212 region of CINP was also generated. Disruption of the interaction was confirmed through affinity pull-down assays (**Figure 5F**).

### The architecture of the 55LCC complex

To characterize the structural organization of the cofactors CINP and C1ORF109 with the AAA+ ATPases SPATA5 and SPATA5L1, we transiently overexpressed all four proteins in HEK293-6E cells using MultiMam^24^ expression vectors. We then affinity-purified complexes containing SPATA5/SPATA5L1 with CINP and C1ORF109, as previously described (**Figure 6A, i, Supplementary Figure S3A-C**). The molecular weight and stoichiometry of the resulting heterooligomers were analyzed by size-exclusion chromatography (SEC) and mass photometry in the absence of nucleotide analogues. These analyses showed that the purified oligomers ranged from ∼ 500 to 1500 kDa under our experimental conditions (**Supplementary Figure S7A**).

**Figure 6.**
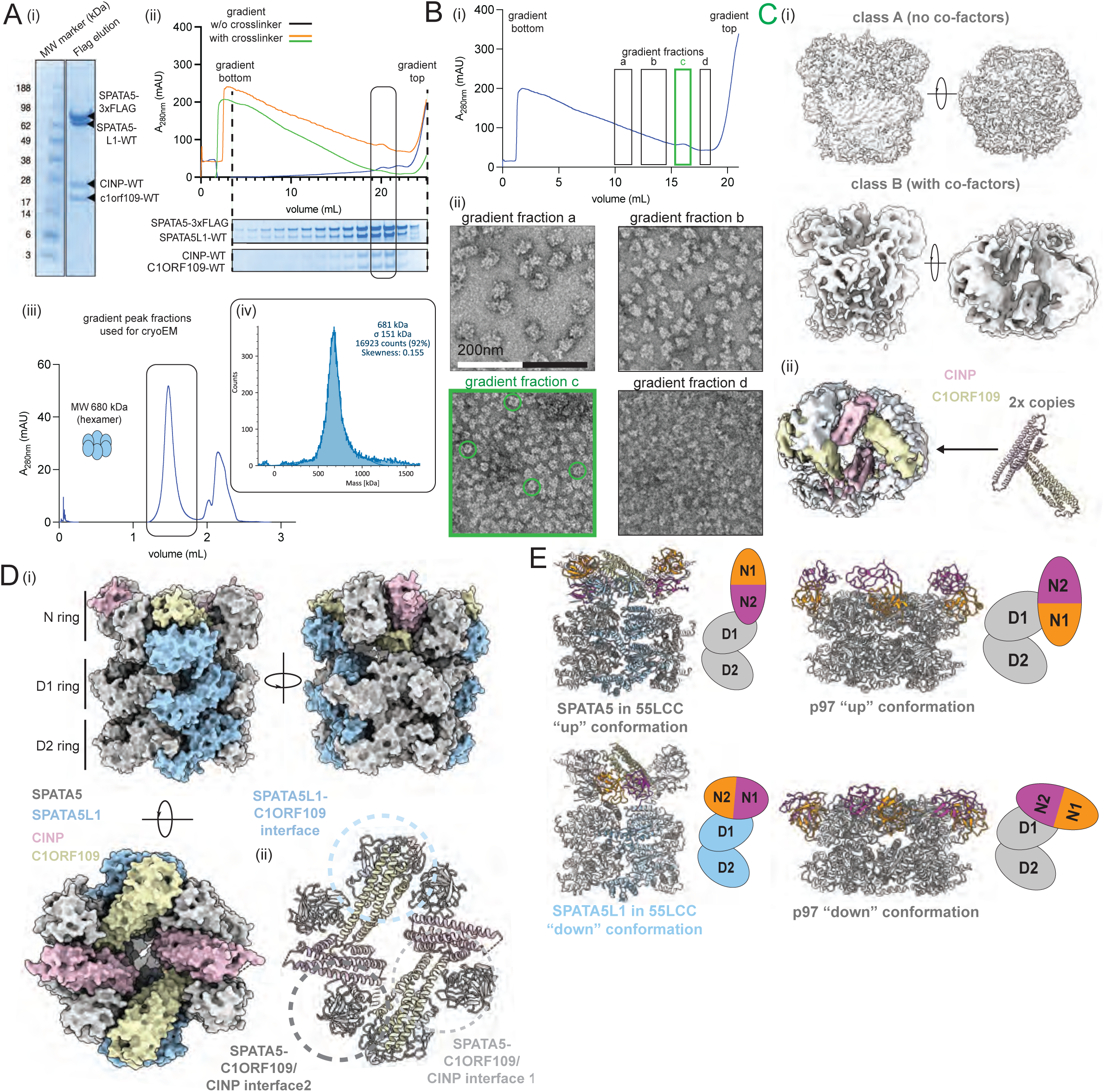
55LCC oligomer stoichiometry and architecture. (**A**) Recombinantly expressed and affinity-purified SPATA5-SPATA5L1-CINP-C1ORF109 (i) is pre-incubated with non-hydrolyzable ATPγS and used as input for sucrose density gradient centrifugation (SDGC) (ii). The four proteins sediment over a broad range of the 10-30% sucrose gradient but are slightly enriched in one peak fraction of distinct molecular weight in the presence and absence of protein crosslinkers. Analysis of the crosslinked SDGC peak fractions by SEC (iii) as well as mass photometry (iv) reveals a stabilized ATPγS-bound oligomeric species of a molecular weight of around 681 kDa ± 20-30 kDa (corresponding to heterohexameric ATPase ring of SPATA5/SPATA5L1 subunits in complex with cofactors CINP/C1ORF109). (**B**) Negative stain TEM analysis of crosslinked SDGC gradient fractions (i) from affinity-purified ATPγS-bound SPATA5-SPATA5L1-CINP-C1ORF109 assemblies. From bottom (30% sucrose) to top (10% sucrose) of the gradient, the four proteins occur within various heterogeneous assemblies (ii): (fraction a) average size ranging from 35-50 nm in diameter (potentially small fraction of contaminant ribosomal subunit-bound species), (fraction b) elongated particles of about 20-40nm length, (fraction c) particles of 20-25 nm in diameter and (fraction d) particles of less < 20nm diameter. Only in the enriched fraction c, it was possible to identify ring-shaped particle images corresponding to hexameric ATPase ring top views (green circles). (**C**) Single-particle cryo-EM analysis of crosslinked fraction c yields two different types of 3D reconstructions after masked classification (i). Both reveal a characteristically shaped hexameric heteromeric AAA+ ATPase ring, with D2, D1 and N-terminal domain rings of SPATA5/SPATA5L1 stacked on top of each other. While N-terminal domains are flexible and badly resolved in map A, extra densities at the N ring in map B could be attributed to two copies of the CINP-C1ORF109 (pink and yellow, respectively) (ii) dimer as determined by X-ray crystallography (**Figure 5**). D 55LCC model created by docking AlphaFold 2 models35,36 of four copies of SPATA5 (grey) and two copies of SPATA5L1 (blue) as well as two CINP-C1ORF109 heterodimers into map B in agreement with PDB 8RHN7. (**E**) Structural comparison of 55LCC complex with different hexameric conformations adopted by AAA+ ATPase p9733. F Verification of cofactor/ATPase interactions by in vitro pull-down experiments pulling on recombinant CINP/C1ORF109 incubated with SPATA5 wild-type (i), N-terminally truncated SPATA5 (N65) (ii) and SPATA5L1 (iii). ^8^

We next optimized purification conditions by testing SEC and sucrose density gradient centrifugation (SDGC) under various salt and nucleotide conditions (ATP, ADP, AMPNP, and ATPγS). SDGC in the presence of non-hydrolyzable ATP analogs – particularly ATPγS – enriched a distinct oligomer population containing all four proteins, with or without glutaraldehyde/BS3 crosslinking (**Figure 6A, ii-iii**). Based on the known 1:1 assembly ratio of CINP and C1ORF109 (**Figure 5**) and a measured molecular weight of ∼ 681 kDa (± 20– 30 kDa) by mass photometry (**Figure 6A, iv**), we inferred that the complex comprises six ATPase subunits (SPATA5/SPATA5L1) along with at least one copy each of CINP and C1ORF109. These findings suggest the formation of a hexameric heterooligomer, likely with a 4:2:2:2 stoichiometry for SPATA5:SPATA5L1:CINP:C1ORF109, in agreement with recently published high-resolution structures of the 55LCC complex^8,10^.

Negative-stain electron microscopy (EM) of crosslinked SDGC fractions confirmed multiple assembly states (**Figure 6B, i**). Fraction (a) contained particles ∼ 35–50 nm in diameter, consistent with pre-ribosomal particles; fraction (b) showed asymmetric particles ∼ 20–40 nm in size, likely representing higher oligomers; fraction (c) contained ∼ 20 nm particles with dimensions consistent with hexameric AAA+ ATPAse assemblies (similar in size to Drg1 hexamer^17^, and the segregase p97^25^); and fraction (d) displayed heterogeneous particles < 20 nm, likely representing dissociated ATPase subunit trimers or dimers (**Figure 6B, ii**). All four proteins were detected across these fractions, but the most homogeneous particle population was found in the enriched peak of fraction (c). Here, we identified ring-shaped particles and conducted negative-stain EM data collection, particle picking, extraction, and 2D classification, which yielded class averages consistent with projection views of AAA+ ATPase hexameric ring assemblies (**Figures 6B, ii, Supplementary Figure S7B**).

Single-particle cryo-EM analysis of this fraction allowed us to distinguish complexes with and without bound cofactors. We reconstructed two 3D maps at ∼12 Å (map A) and ∼15 Å (map B) resolution (**Figure 6C, i and Supplementary Figure S7C–D**). Both maps displayed the characteristic AAA+ ATPase hexamer canonical architecture, with three stacked rings corresponding to the D2, D1, and N-terminal domains (**Figure 6C**). An AlphaFold-predicted^26,27^ SPATA5:SPATA5L1 4:2 heterohexamer model (**Supplementary Figure S7E**), consistent with the observed stoichiometry and molecular weight, docked well into the D1/D2 regions of both maps. In map A, the N-terminal domains were poorly resolved, likely due to conformational flexibility, suggesting this map represents the ATPase complex alone (55L) (**Figure 6C, i**). In contrast, map B showed additional density above the N-terminal domain ring, allowing placement of the CINP–C1ORF109 heterodimers based on its crystal structure (**Figure 5, 6C, ii, Supplementary Figure S7F**), resulting in a reconstructed 55LCC complex resembling the recently published structures^8,10^ (**Figure 6D**). Notably, the linkers connecting the D1-D2 and D1-N-terminal domains were not resolved in the EM density, likely due to their flexibility, consistent with AlphaFold-MultiMer predictions^26,27^ (**Supplementary Figure S7E**).

Our low-resolution cryo-EM reconstructions complement recent high-resolution structures of the 55LCC complex, including a 4.5 Å structure from recombinant insect cell expression^8^ and a 3.8 Å structure from human cells expressing the stabilized SPATA5 2EQ mutant^10^. While these studies achieved superior resolution, enabling detailed atomic modeling, our analysis of wild-type protein expressed in mammalian cells without stabilizing mutations shows the native conformational heterogeneity of the complex, leading to a limited resolution. The lack of defined density on the D1-D2 and D1-N-terminal domain linker regions reflects intrinsic dynamics in the wild-type complex^8,10^.

In the cryo-EM structure of the 55LCC complex, each SPATA5 N-terminal domain adopts an ‘up’ conformation at the interface between a CINP and a C1ORF109 molecule. Despite its distinct heterohexameric 4:2 composition, 55LCC exhibits N-terminal domain movements similar to the nucleotide-driven conformational changes observed in its functional and structural homologs such as Drg1 and p97^17,28^, though in 55LCC, this flexibility seem to be additionally coupled to cofactor binding rather than being purely nucleotide-dependent **(Figure 6E**). This ‘up’ orientation enables the SPATA5 N-terminal domain to engage both cofactors simultaneously through two distinct surfaces enriched in conserved residues (**Supplementary Figure S8–S10**). In contrast, the SPATA5L1 N-terminal domains are positioned closer to their respective D1 domains, rather than aligning horizontally with the SPATA5 N-terminal domains. While they do not fully adopt the canonical ‘down’ conformation seen in ADP-bound p97 hexamers, their placement is stabilized through extensive interactions with C1ORF109 (**Figure 6D, ii**). Notably, SPATA5L1 shows little to no direct interface with CINP (**Figure 6D, ii).**

### The ATPase activities of SPATA5 and SPATA5L1 are required for 55LCC function

To characterize the relevance of the ATPase function of SPATA5 and SPATA5L1, and the relevance of specific interactions identified within the 55LCC complex, we developed a system to acutely swap endogenously expressed SPATA5, SPATA5L1, CINP, or C1ORF109 with wild-type or mutant transgenes of the respective proteins. For this, we used SPATA5^dTAG^, SPATA5L1^dTAG^, CINP^dTAG,^ and C1ORF109^dTAG^ expressing cells, and in this genetic background, wild-type or mutant transgenes were ectopically expressed by stably introducing the respective transgenes fused with HALO-Tag via CRISPR in the AAVS1 safe-harbor locus. HALO-Tag fusion allows acute transgene depletion using the HaloPROTAC-E^29^. In this system, treatment with dTag^V^-1 induces the depletion of the endogenously expressed 55LCC constituents, while HaloPROTAC-E induces degradation of the ectopically expressed proteins, and combined treatment with dTAG^V^-1 and HaloPROTAC-E causes degradation of both the endogenously expressed and ectopically expressed proteins (**Figure 7A**).

**Figure 7.**
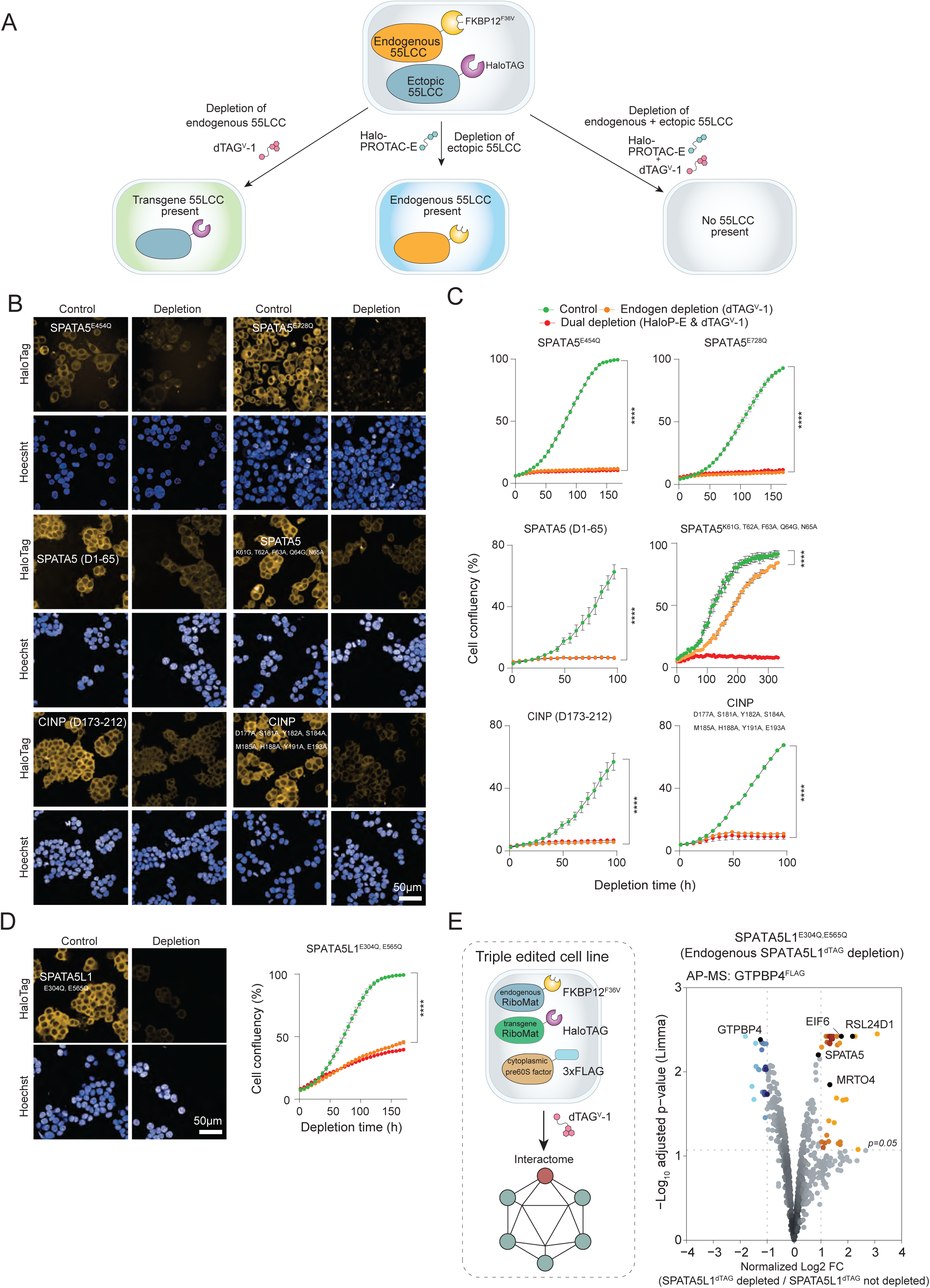
Requirement of SPATA5 and SPATA5L1 ATPase activities and specific interaction regions for the 55LCC function. (**A**) Strategy for acute swapping of endogenously expressed 55LCC constituents with their respective ectopically expressed counterparts. Endogenously expressed 55LCC constituents were fused with dTAG (**Figure 3A**), and in these backgrounds, their respective counterparts were expressed ectopically via targeted integration of transgene expression cassettes into the AAVS1 safe-harbor locus. Ectopically expressed proteins were fused with HaloTAG. The fusion proteins can be depleted in a desired combination by treating the cells with dTAG^V^-1 and HaloPROTAC-E. (**B**) Micrographs showing ectopic expression of the indicated transgenes, and their acute depletion by HaloPROTAC-E (250 nM). (**C**) Growth curves of cell lines expressing the indicated transgenes after the specified treatments (*n*=3, Mean ± SEM). (**D**) Micrographs (left panel) and growth curves (right panel) of cell line expressing SPATA5L1^E340Q,E565Q^ after the indicated treatments (*n*=3). (**E**) Schematic of the experimental setup using SPATA5L1 dual ATPase-dead variants and additional endogenous 3xFLAG-tagging of GTPBP4, enabling comparative AP-MS analysis of GTPBP4 interactomes in the presence of wild-type versus mutant SPATA5L1 (E304Q, E565Q) protein variants. Volcano plot of GTPBP4-FLAG AP-MS results comparing the enrichment at GTPBP4 in the presence of SPATA5L1 ATPase-dead variant versus endogenous SPATA5L1, highlighting differences in GTPBP4-associated interactions (*n*=3). Statistical significance intervals indicated as: * p<0.05, ** p<0.01, *** p<0.001, **** p<0.0001.

To validate this system, we ectopically expressed wild-type versions for each of the 55LCC constituents in cells in which the respective endogenously expressed 55LCC constituent was fused with dTAG. Expression and HaloPROTAC-E-induced depletion of transgenes verified by immunostaining against HALO-Tag (**Supplementary Figure S11A**). Next, we tested the functionality of the ectopically expressed proteins. For all 55LCC constituents, ectopic expression of the respective wild-type protein rescued the cell proliferation defect caused by depletion of the corresponding endogenous counterpart, while cell proliferation was confirmatively reduced after combined depletion of endogenous and ectopic protein versions (**Supplementary Figure S11B**).

Having demonstrated the suitability of the swapping approach, we introduced various 55LCC mutants to test their functionality. Mutations were introduced to individually ablate the activities of the ATPase domains D1 (SPATA5^E454Q^; SPATA5L1^E304Q^) or D2 (SPATA5^E728Q^; SPATA5L1^E565Q^)^30,31^. Then, we swapped the endogenous SPATA5^dTAG^ and SPATA5L1^dTAG^ with the ATPase dead mutants. For SPATA5, both D1 and D2 ATPase dead mutants failed to rescue the cell proliferation defect caused by endogenous SPATA5^dTAG^ depletion (**Figure 7B-C**). In contrast, mutation in the D1 domain of SPATA5L1 only weakly reduced its ability to rescue cell proliferation, while mutation of the D2 domain fully rescued cell proliferation (**Supplementary Figure S11C-D**). Functional complementation by SPATA5L1 single ATPase dead mutants prompted us to examine the requirements of their combined activities. We mutated both D1 and D2 domains in SPATA5L1 simultaneously and tested their effect on the cell proliferation rescue assay. In this case, SPATA5L1 D1+D2 double ATPase-dead variant failed to rescue cell proliferation (**Figure 7D**).

To confirm that the inability of the ATPase-defective SPATA5L1 to rescue cell proliferation is likely due to its inability to rescue the pre-60S maturation defect caused by the depletion of endogenous SPATA5L1^dTAG^, we generated a triple-edited cell line. In the SPATA5L1^dTAG^

/ SPATA5L1^HALO_D1+D2_dead^ background, the endogenous GTPBP4 was fused with 3xFLAG (**Figure 7E**). Then, we compared the GTPBP4 interactomes, with and without depletion of the endogenous SPATA5L1^dTAG^. This revealed increased enrichment of pre-60S-associated RBFs, such as EIF6, MRTO4, RSL24D1, as well as SPATA5. This suggests that the ATPase-dead SPATA5L1 fails to promote the release of pre-60S-associated factors while 55LCC remains intact and associated with the pre-60S.

Together, these results demonstrate the essentiality of both ATPase domains of SPATA5 and indicate that each domain has an individual, non-redundant function in supporting cell growth. In contrast, the activity of individual SPATA5L1 ATPase domains appears less critical, but their combined activities are required for supporting pre-60S maturation and cell proliferation, suggesting a potentially redundant function between these two domains.

### CINP and C1ORF109 heterodimerization is required for 55LCC function

Using the above-described transgene complementation approach, we tested the requirement of key interactions in 55LCC identified in our structural analyses (**Figures 7B-C**). Ectopic expression of SPATA5 lacking its N-terminal region (1–65), which is posited to interact with CINP–C1ORF109, failed to rescue proliferation in endogenous SPATA5^dTAG^-depleted cells. A SPATA5 with five amino acid mutations (N61G, T62A, F63A, Q64G, N65A) in this region partially rescued proliferation, suggesting reduced interaction efficiency.

To further investigate whether the individual interaction of CINP with SPATA5 was sufficient for 55LCC function or the binding of the stable coiled-coil CINP-C1ORF109 heterodimer is required (**Figure 5**), we depleted endogenous CINP^dTAG^ and expressed CINP mutants that fail to interact with C1ORF109 (**Figures 7B-C**). Neither mutant rescued proliferation in endogenous CINP^dTAG^-depleted cells. These findings support that CINP– C1ORF109 dimerization is essential for 55LCC function.

These results show that the 55LCC complex’ function in ribosomal biogenesis depends not only on the ATPase activities of SPATA5 and SPATA5L1, but also on the interactions with the two coiled-coil cofactors.

## Discussion

The 55LCC complex characterized in our study supports the increased complexity of ribosome biogenesis not found in lower eukaryotes. Our work confirms the recently reported structure and function of the 55LCC complex in cytoplasmic pre-60S assembly^8–10^.

### Metazoan-specific 55LCC regulates pre-60S assembly

Our AP-MS and functional analyses show that 55LCC is essential for the cytoplasmic maturation of pre-60S. This suggests 55LCC serves as a function analogous to yeast Drg1^19^. Notably, the activities of both ATPase domains are essential for SPATA5 function. In contrast, in SPATA5L1, activities of individual ATPase domains appear dispensable, but their combined ATPase activities are required to support cell proliferation. This suggests that SPATA5 activities are more crucial than those of SPATA5L1.

Mutations in multiple RBFs are also associated with neuronal developmental disorders^32^. Mutations in human SPATA5 and SPATA5L1 are associated with reduced global growth, developmental delay, microcephaly, and intellectual impairment^33,34^. It is tempting to postulate that developmental delay, microcephaly, and intellectual impairment in SPATA5 mutant patients may be due to impairment in ribosome biogenesis.

### Does 55LCC have roles beyond pre-60S assembly?

Our biochemical and cell biological analyses unequivocally demonstrate that 55LCC plays a role analogous to yeast Drg1 in the cytoplasmic maturation of pre-60S^19^. However, single-celled organisms lack SPATA5L1, CINP, and C1ORF109, and yeast Drg1 alone is sufficient for this process. Why eumetazoans additionally require SPATA5L1, CINP, and C1ORF109 remains a mystery. One possibility is that 55LCC evolved additional regulatory functions beyond ribosome biogenesis.

During the course of our work, Krishnamoorthy et al. reported that 55LCC interacts with replisome components and 55LCC-mediated proteostasis is important for replication fork progression and genome stability^8^. In our extensive interaction analyses of endogenously engineered 55LCC components, we consistently found interaction of 55LCC with RPs and RBFs. However, we found no reproducible interactions with proteins involved in DNA replication and the cell cycle. Furthermore, in our subcellular localization analyses, performed on endogenously expressed proteins, all four members of 55LCC are almost exclusively found in the cytoplasm. The reason for this discrepancy remains unclear. We cannot rule out the possibility that a small amount of 55LCC is present on the chromatin and interacts with proteins involved in regulating DNA replication and cell cycle, but the enrichment conditions in our analyses may be too harsh to detect weak or transient interactions.

### Insights into the 55LCC complex structure

The 55LCC complex exhibits substantial conformational flexibility, particularly in the N-terminal domains of the ATPases, which is modulated by nucleotide state and cofactor binding. This dynamic architecture likely contributes to the difficulty in isolating a well-defined oligomeric assembly. Such flexibility is consistent with observations from p97 structural studies, where the D1-N-terminal domain linker remains mobile in most nucleotide states except during ATP hydrolysis transition^35^.

In our 55LCC preparations, ATPγS stabilizes the complex, suggesting that nucleotide binding is essential for conformational priming rather than hydrolysis itself. This supports the idea that cofactor binding may preferentially occur in ATP-bound states^36^ (**Figure 6**), further reinforcing the parallels with p97. Consistent with our cryo-EM data, both Krishnamoorthy et al. and Dai et al. reported substantial flexibility at the N-terminal domain and ATPase rings interfaces^8,10^. Indeed all published structures were obtained in the presence of ATP or ATPγS to lock the complex in defined conformational states^8,10^. In agreement with these observations, SPATA5/SPATA5L1 ATPase-deficient mutants fail to yield stable complexes in our purifications (**Supplementary Figure S3C**). However, Dai et al. successfully determined a structure with SPATA5 ATPase mutants in the Walker B motif^10^, indicating that these variants can be stabilized when bound to the pre-60S substrates.

Conversely, SPATA5L1 N-terminal domains adopt a distinct position compared to SPATA5 N-terminal domains, residing in a lower horizontal plane within the complex and interacting with the cofactors from a different angle (**Figure 6E**). Both studies confirm this orientation difference^8,10^. In our structures at low resolution, it remains uncertain whether SPATA5L1 subunits are nucleotide-bound, particularly as activities of individual SPATA5L1 ATPase domains do not appear to be a critical requirement for 55LCC integrity and function (**Figure S11D**). The specific ATP and cofactor-induced conformation would indicate an explanation for the heterogeneity of the 55LCC oligomers, as after ATP binding, the complex might only exist in a transient non-target-bound state.

Although proteomic analyses (**Figure 2A**) revealed enrichment of ribosomal proteins and maturation factors within the SPATA5/SPATA5L1 interactomes, direct biochemical evidence for stable interaction with pre-60S subunits remained limited in our study. We hypothesized a transient association of 55LCC complex with late-stage pre-60S particles through cofactor-mediated recognition, potentially involving the GTPBP4^37^. This prediction was directly confirmed by Dai et al., who demonstrated that CINP interacts directly with GTPBP4, forming the first interface between 55LCC and pre-60S^10^. The structure also revealed a second interface between CINP/SPATA5 and ES27A, and showed that PA2G4 (human Arx1 homolog) stabilizes ES27A conformation to indirectly facilitate 55LCC binding, rather than directly interacting with the 55LCC complex^10^. In yeast, Rlp24 recruits Drg1 onto the ribosome through its C-terminal 53 amino acids^38^, which show no sequence similarity with RSL24D1, suggesting distinct interaction modes. However, Dai et al. indicate that the fundamental principle of RSL24D1 C-terminal mediated recruitment is conserved: based on their partially resolved 55LCC:pre-60S structure, the unresolved C-terminal tail of RSL24D1 seems to be at the correct distance to reach 55LCC ring pore^10^, despite the lack of sequence conservation. This arrangement supports our hypothesis that ATP-driven conformational changes within the 55LCC complex could promote coordinated release of GTPBP4 and RSL24D1 maturation factors, which have been proposed to dissociate together during cytoplasmic pre-60S remodeling^19,38^, in agreement with our previous observations (**Figure 4, Supplementary Figure S4 and S5**).

This model aligns with known functions of other AAA+ ATPases like p97 and Rix7, which operate as nucleotide-dependent remodeling machines in processes such as ER-associated degradation and nucleolar pre-ribosome disassembly^39,40^. Like these complexes, 55LCC may act as a regulated, substrate-specific disassembly factor, engaging its targets only under controlled conformational or temporal conditions. The inability to capture stable interaction intermediates could thus reflect a combination of rapid turnover, substrate specificity, and dependence on yet-unidentified adaptor proteins or structural cues.

## Methods

### Cell culture

Human HCT116 and HEK-293-6E cells were purchased from American Type Culture Collection (ATCC) and National Research Council Canada, respectively, and cultured in DMEM GlutaMAX media (Gibco). Media was supplemented with 10% fetal bovine serum (FBS, Sigma-Aldrich) and 1x Penicillin-Streptomycin and 2 mM L-Glutamine (Gibco). For SILAC^41^ (Stable Isotope Labeling with Amino Acids in Cell Culture) experiments, cells were cultured in customized DMEM (C.C.Pro GmbH) without L-Lysine; L-Arginine, L-Glutamine supplemented. Medium was supplemented with 1x Penicillin-Streptomycin, 200 nM L-Glutamine (Gibco), and 10% dialyzed FBS (Sigma-Aldrich). Respective labeled L-Lysine HCL (84 mg/mL; Cambridge Isotope Laboratories Inc.) and L-Arginine HCL (146 mg/mL; Cambridge Isotope Laboratories Inc.) were added as well as 10% dialyzed FBS (Sigma-Aldrich), 1x Penicillin-Streptomycin and 2mM L-Glutamine (Gibco). For SILAC experiments, cells were labeled for three weeks with SILAC medium supplemented with SILAC amino acids of the following isotopic compositions before respective experiments: ‘Light’ L-Arginine-^12^C_6_^1^H ^14^N ^16^O_2_, ‘Light’ L-Lysine-^12^C_6_^1^H ^14^N ^16^O_2_, ‘Medium’ L-Arginine-^13^C_6_^1^H ^14^N ^16^O_2_, ‘Medium’ L-Lysine-^12^C_6_^1^H_14_ ^2^H ^14^N ^16^O_2_, ‘Heavy’ L-Arginine-^13^C_6_^1^H_14_ ^15^N_2_^16^O_2_, ‘Heavy’ L-Lysine-^13^C_6_^1^H_14_ ^15^N ^16^O_2_. Cells were grown at 37°C in a humidified incubator at 5% CO_2_ and passaged at 70-80% confluence. For assessments, cells were seeded with assay-specific densities.

### Generation of cell lines

Cell line engineering was performed using CRISPR/Cas9 and separate donor and Cas9 plasmids targeted to specific genomic sites. The plasmids used in this study are listed along with the resulting engineered cell lines (**Supplementary Table 1**). Homology arms (HA) were primarily amplified from genomic DNA, and in some cases, synthesized (GeneArt Strings). A multiple cloning site (MCS) was inserted in between HAs in-frame with the targeted Exon using restriction enzyme cloning or Gibson Assembly. Fusion tags contained a drug resistance selection gene (puromycin: *pac* from *Streptomyces alboniger*, hygromycin: *hph* from *Escherichia coli,* nourseothricin: *nat* from *Streptomyces noursei*, or blasticidin: *bsr* from *Bacillus cereus*), mostly separated by a P2A self-cleavage site from the fusion protein/tag (**Supplementary Table 1**). Guide RNA sequences were designed with the help of CRISPOR^42^ and inserted into the vector px330 encoding Cas9. For the editing, cells were seeded in a 6-well dish (300,000 cells/well) and transfected ∼24 h after seeding. Cas9-endonuclease and HDR donor plasmids were simultaneously introduced using TurboFect Transfection Reagent following the manufacturer’s instructions (Thermo Fisher Scientific). The transfection mix was removed after incubation and an additional ∼24h of recovery, cells were split, and edited cells were selected based on the introduced antibiotic selection marker. The resulting cell colonies were clonally purified, and editing success was screened using oligonucleotides binding to DNA regions flanking the HAs (**Supplementary Table 1**). Clones with the desired genotype were confirmed, expanded, characterized, stocked, and used for experiments.

For initial interactome screenings and monitoring of endogenous protein distribution, we tagged POIs with GFP, mCherry, or 3xFLAG at indicated termini. Cell lines for acute protein depletion were generated by tagging the 55LCC constituents with FKBP12F36V (dTAG)^20^ or green fluorescent protein-FKBP12F36V (GFP-dTAG).

### Generation of stable transgene-expressing cell lines

mRNA was extracted from HCT116 cells using the RNeasy Kit (Qiagen), and cDNA was synthesized with SuperScript™ IV Reverse Transcriptase (Invitrogen), following the manufacturer’s instructions. The native coding sequences (CDSs) of the proteins of interest were amplified from the resulting cDNA using custom-designed oligonucleotides (**Supplementary Table 1**). CDSs were cloned into an AAVS1-integrating donor vector, where specified, desired mutations were introduced and confirmed by sequencing. When successfully integrated, the vector expresses desired POI variants under the control of an hPGK promoter, while all variants were fused to HaloTag and linked to an antibiotic resistance gene via a P2A for the selection of edited cells. Before testing the mutated protein variants, the system’s functionality was confirmed through the successful rescue of the endogenously 55LCC constituent-depleted cell lines using the respective wild-type transgene. Integration of the transgenes into the AAVS1 locus was validated by genomic PCR using homology arm-flanking oligonucleotides, and the correctness of variants was confirmed by Sanger sequencing. Halo-Click staining (Janelia Fluor® 549 HaloTag® Ligand) was used to confirm correct subcellular localization and HaloP-E-induced depletion of the transgene-expressed proteins.

### Immunofluorescence cytochemistry

For immunocytochemistry (ICC) experiments, cells were seeded at a density of 15,000 cells per well on Perkin Elmer 96-well glass-bottom plates (Cell Carrier-96 Ultra), which had been pre-coated with 50 mg/mL Poly-L-Lysine diluted in 1x PBS, for 1 h at 37°C. After 24 h of attachment time, assays were started, and at the end of treatment time, cells were fixed with 3.7% formaldehyde in 1x PBS for 15 min at RT. Following fixation, the cells were washed three times with 1x PBS, permeabilized with 0.5% Triton X-100 in 1x PBS for 10 min, and blocked with 5% FBS in 1x PBS for 1 h at RT. Cells were incubated with primary antibodies (Anit-GTPBP4 (1:100), Abcam ab184124; Anti-RSL24D1 (1:200), Sigma-Aldrich (HPA062724); Anit-MRTO4/Anti-C1orf33 (1:100), Sigma-Aldrich (WH0051154M1); Anit-GFP antibody rabbit polyclonal [PABG1] (1:250), Chromotek/Proteintech (PABG1-100); 5f8-α-Red FPs (RFP) (1:250), Chromotek/Proteintech (5f8); ANTI-FLAG® M2 antibody (1:2,500), Merck (F3165-1MG) diluted in 2% BSA in 1x PBS for 1 h at RT or overnight at 4 °C. After three additional PBS washes, cells were incubated with Alexa Fluor conjugated second ary antibodies (1:1,000, Thermo Fisher Scientific, A28175, A11036, A11006) and nuclei were counterstained with Hoechst 33342 (1:10,000) for 1h. Finally, cells were washed three times with 1x PBS and remained in 1x PBS for Microscopy. For Halo-Tag visualization, living cells were incubated with Janelia Fluor® 549 HaloTag® Ligand (Promega, 1:1,000) and Hoechst 33342 (1:10,000) added to Gibco™ FluoroBrite™ DMEM medium for 20min. After three 1xPBS washes, cells were left in FluoroBrite medium for a 20-minute wash-out in the incubator. Finally, cells were washed three times with 1x PBS and were directly imaged.

### High-throughput microscopy

Fluorescence images were captured using a PerkinElmer Phenix spinning disk microscope, equipped with two 16-bit sCMOS cameras (2 160×2,160 pixels, 6.5 µm pixel size). Depending on the experiments, different objectives were used: a 40x objective (NA 1.1 W Plan Apochromat/WD 0.62 mm) or a 63x objective (NA 1.15 W Plan Apochromat/WD 0.6 mm). Different fluorescence channels were acquired separately. To ensure consistency and comparability, identical settings were applied within each experiment or antibody setup. Each experiment acquisition was performed in triplicate, several images were acquired per technical replicate (>10) in an unbiased and automated manner, and the resulting images were quantified as described in the data analysis section.

### Bright-field image-based cell proliferation analyses

Growth assessment was conducted using the Incucyte Live-Cell Analysis System (Sartorius). Bright-field images were captured at 10x magnification in a time series, with multiple images taken for each condition and replicate. Imaging data were processed using the manufacturer’s software (Incucyte 2022B Rev2), utilizing preset parameters optimized for cell type recognition (classic confluence, minimum area 200 μm²). Cells were seeded one day before the experiment, and the medium was replaced with the indicated treatments. A 2h lag phase was incorporated at the start of the assessment to prevent condensation on the plates. The resulting mean percentages of area coverage, quantified for each image per condition (well), were plotted.

### Viability Assessment via Tryptophan Staining

Cells were seeded in 12-well plates at a density of 30,000 cells per well and treated from the time of seeding. At the indicated time points, media was collected in a tube, and the wells were rinsed with 200 µL of 1× PBS, which was also collected. Subsequently, 150 µL of 0.05% trypsin-EDTA (Thermo Fisher Scientific) was added to each well and incubated for 4 minutes at 37 °C to detach the cells. The previously collected medium was then used to resuspend and transfer the cells. Cell suspensions were centrifuged for 3 minutes at 600 rcf, and the supernatant was carefully aspirated. Resulting pellets were resuspended, mixed thoroughly, and each suspension was mixed 1:1 with a tryptophan-based viability stain (Thermo Fisher Scientific) following the manufacturer’s instructions. The stained samples were loaded into Countess™ cell counting chamber slides and analyzed using the Countess™ II FL Automated Cell Counter (Thermo Fisher Scientific), using standardized settings. The instrument quantified total cell number and viability percentage based on dye exclusion/inclusion, allowing discrimination between live and dead cells. For each cell line and time point, samples of one cell type were processed and measured sequentially, with three technical replicates analyzed per condition before proceeding to the next replicate set.

### Cloning and expression of proteins in mammalian cells

Two types of open reading frames (ORFs) were used for transient overexpression in HEK293-6E cells. Commercially synthesized (Genscript) codon-optimized sequences for SPATA5, SPATA5L1, CINP, and C1ORF109 were initially transferred into insect cell MultiBac^43^ overexpression vectors. SPATA5 and C1ORF109 sequences were inserted into multiple cloning sites 2 and 1 of the pFL acceptor vector (using restriction enzymes XmaI/NheI and EcoRI/BamHI1, respectively), while SPATA5L1 and CINP were incorporated into multiple cloning sites 1 and 2 of donor vector pSL (using restriction enzymes RsrII/SaII, as well as NheI/XmaI). From those dual plasmids, a quadruple MultiBac co-overexpression vector was created using the Cre-LoxP system and Blue/White clone selection according to the MultiBac manual. For expression in HEK293-6E cells, the resulting ORFs were later transferred into mammalian MultiMam^44^ overexpression acceptor vector pACEMam1 (SPATA5 or SPATA5L1 for individual overexpression), and donor vectors pMDC (SPATA5L1), pMDS (CINP), and pMDK (C1ORF109) using PCR-based In-Fusion cloning (Takara Bio). For inserting a C-terminal TEV-cleavable 3xFLAG tag instead of the original SPATA5 stop codon and for creating the SPATA5 N-terminal truncation mutant N65 (comprising only residues 66-893 of SPATA5 WT) In-Fusion cloning (Takara Bio) was used. ATPase deficiency point mutations for SPATA5 (D1 – E454Q and/or D2 – E728Q), as well as SPATA5L1 (D1 – E304Q and/or D2 – E565Q), were introduced in the above vectors using site-directed mutagenesis. Again, plasmids were combined into one quadruple MultiMam co-overexpression vector using Cre recombinase (New England Biolabs). In a third round, to test the possible effects of the original codon sequence on protein expression and protein stability, the original CDS sequences (prepared as described in the preceding Methods section Generation of stable transgene cell lines) were inserted into the existing MultiMam plasmids instead of the codon-optimized sequences, again using In-Fusion cloning (Takara Bio). This third set of expression vectors was used for expressing and purifying proteins used in cryo-EM and *in vitro* pull-down assays.

### Cloning and expression of native proteins in bacterial cells

Proteins were expressed with an N-terminal hexahistidine (His)-tag and a TEV cleavage site followed by the full-length CINP (M1-K212), or truncated ΔCINP (V18-K212) and full-length C1ORF109 (M1-E203), respectively. A truncated ΔCINP (V18-K212)-C1ORF109 construct was designed to facilitate complex crystallization due to the flexibility of the N-terminal end of CINP. For this, commercially synthesized and codon-optimized CINP and C1ORF109 coding sequences (Genscript) were cloned either alone (for single expression constructs) or together (for double co-expression constructs) into multiple cloning site 1 (CINP, using restriction enzymes *BamHI* and *NotI* (New England Biolabs)) and 2 (C1ORF109, using restriction enzymes *NdeI* and *XhoI* (New England Biolabs)) of the pETDuet expression vector (Novagen). The N-terminally truncated version of ΔCINP (V18-K212) was created by a PCR-based deletion using In-Fusion cloning (Takara Bio).

Proteins were expressed in *E. coli* BL21 pRARE cells (Thermo Fisher Scientific). *E. coli* cultures were grown at 37 °C in Lysogeny Broth (LB) medium with 100 mg/L ampicillin to an OD600 of 0.6. Overexpression of proteins was induced with 0.5 mM IPTG and grown overnight at 18 °C. Cells were harvested by centrifugation at 4,000 rcf (F9S-4×1000y rotor, Thermo Fisher Scientific) for 15 min, flash-frozen in liquid nitrogen, and stored at −80 °C.

### Expression of selenomethionine-incorporated proteins in bacteria

For the expression of selenomethionine-incorporated proteins, the same DNA constructs have been used. Proteins were expressed in *E. coli* BL21 pRARE cells (Thermo Fisher Scientific). *E. coli* cultures were grown at 37 °C in SelenoMethionine Medium Complete (Molecular Dimensions) with 100 mg/L ampicillin to an OD600 of 0.6. Overexpression of proteins was induced with 0.5 mM IPTG and grown overnight at 18 °C. Cells were harvested by centrifugation at 4,000 rcf (F9S-4×1000y rotor, Thermo Fisher Scientific) for 15 min, flash-frozen in liquid nitrogen, and stored at −80 °C.

### Purification of 55LCC and SPATA5 variants

Cell pellets were resuspended in 100 mL per 30 g pellet in ice-cold lysis buffer (100 mM HEPES-KOH pH 7.5, 250 mM NaCl, 2 mM MgCl_2_, 10% glycerol, 0.008% NP-40, 0.25% Tween20, 4 tablets of cOmplete Inhibitor cocktail EDTA Free (Roche) per 250 mL, benzonase 8 uL per 250 mL). The resuspended pellets were stirred for 20 min at 4 °C. Lysis was completed by manual homogenization in a dounce tissue grinder (DWK Life Sciences) and sonication on ice using an Ultrasonic Liquid Processor (15 s ON/20s OFF, 40% amplitude, for 4 min, Misonix). The cell lysate was cleared by centrifugation at 200,000 rcf for 1 h at 4 °C (45 Ti rotor, Beckman), and the soluble fraction was passed through a 0.45 µm vacuum filter (CELLTREAT). The soluble fraction was loaded into an Econo-Pac chromatography column (BioRad) packed with 1.5 mL ANTI-FLAG M2 affinity gel (Millipore) equilibrated in FLAG buffer (100 mM HEPES-KOH pH 7.5, 250 mM NaCl, 2 mM MgCl_2_, 10% glycerol). The sample was washed with 35 column volumes (CV) of FLAG buffer, then 25 CV of FLAG buffer supplemented with 5 mM ATP magnesium salt (Sigma-Aldrich), followed by a wash with another 35 CV of unsupplemented FLAG buffer. Bound proteins were eluted by five subsequent washes (0.7 CV, 30 min incubations) with 0.4 mg/mL 3xFLAG peptide (APExBIO) in FLAG peptide buffer (50 mM HEPES-KOH pH 7.5, 150 NaCl, 150 KCl, 5 mM MgCl_2_). Enriched protein fractions were pooled and concentrated to 5-12 mg/mL, depending on the subsequent use.

### Purification of CINP and ΔCINP-C1ORF109

Cell pellets were resuspended in 30 mL per 5 g pellet in ice-cold lysis buffer (50 mM Tris-HCl pH 8.0, 250 mM NaCl, 2 mM MgCl_2_, 1 mM TCEP, 10 mM imidazole, 0.25% Tween20, 2 tablets of cOmplete Inhibitor cocktail EDTA Free (Roche) per 100 mL, benzonase 3.5 µL per 100 mL). The resuspended pellets were stirred for 1 h at 4 °C. Lysis was completed by manual homogenization in a dounce tissue grinder (DWK Life Sciences) and sonication on ice using Ultrasonic Liquid Processor (20 s ON/20s OFF, 40% amplitude, for 5 min, Misonix). The cell lysate was cleared by centrifugation at 200,000 rcf for 1 h at 4 °C (45 Ti rotor, Beckman), and the soluble fraction was passed through a 0.45 μm vacuum filter (CELLTREAT). The soluble fraction was loaded into a 5 mL HisTrap FF Crude column (Cytiva) equilibrated in buffer HisTrap-A (50 mM Tris-HCl pH 8.0, 150 mM NaCl, 2 mM MgCl_2_, 1 mM TCEP, 20 mM imidazole), and bound proteins were eluted with a linear gradient of 0–100% buffer HisTrap-B (50 mM Tris-HCl pH 8.0, 150 mM NaCl, 2 mM MgCl_2_, 1 mM TCEP, 500 mM imidazole). Enriched protein fractions were pooled together, and His-tag was cleaved overnight with TEV protease using 1:100 TEV:protein ratio. Next, the imidazole concentration was reduced to 25 mM using buffer HiTrap-A (50 mM Tris-HCl pH 8.0, 100 mM NaCl, 2 mM MgCl_2_, 1 mM TCEP). The sample was applied onto a tandem of HisTrap FF crude and HiTrap Q FF column (Cytiva) equilibrated with buffer HiTrap-A (50 mM Tris-HCl pH 8.0, 100 mM NaCl, 2 mM MgCl_2_, 1 mM TCEP) to trap cleaved His-tag and further purify the protein. The protein was eluted with a linear gradient of 0–100% buffer HiTrap-B (50 mM Tris-HCl pH 8.0, 1 M NaCl, 2 mM MgCl_2_, 1 mM TCEP). Then, the protein was concentrated and further purified by SEC using a Superdex 75 10/300 column (Cytiva) equilibrated in Crystal buffer (100 mM Tris-HCl pH 7.5, 100 mM NaCl, 1 mM TCEP). Fractions containing pure protein were pooled, concentrated to 9.2 mg/mL (for CINP), 13.5 mg/mL (for native ΔCINP-C1ORF109), 13.6 mg/mL (for selenomethionine-incorporated ΔCINP-C1ORF109), flash-frozen in liquid nitrogen, and stored at −80 °C.

### Crystallization

Experiments were set up using the sitting-drop vapor-diffusion method. Reservoir solution (0.236 M sodium citrate pH 7.0, 23.4% 2-propanol, 0.1 sodium cacodylate pH 7.0) was dispensed using the Dragonfly Crystal robot (sptlabtech), and the drops were dispensed using the Mosquito LV Crystal robot (sptlabtech) in the following proportions: 500 nL of protein solution (selenomethionine-incorporated ΔCINP-C1ORF109 at 13.6 mg/mL in 100 mM Tris-HCl pH 7.5, 100 mM NaCl, 1 mM TCEP) and 500 nL of reservoir solution. The crystallization trays (sptlabtech) were stored in Rock Imager 1000 (Formulatrix) at 4 °C and checked regularly. Crystals appeared after three days. To freeze the crystals, crystallization trays were first placed on dry ice to promote crystal detachment from the bottom of the well, and glycerol was added to the drop to a final concentration of 20%. Crystals were mounted on loops (Hampton Research) and flash-frozen and stored in liquid nitrogen.

### X-ray data collection, data processing, atomic model building, and refinement

SAD data was collected at the selenium peak wavelength from a single selenomethionine-enriched crystal at the Swiss Light Source synchrotron (Paul Scherrer Institute, Switzerland). Diffraction data processing and preliminary structure solution were conducted using the Crank2 pipeline^45^, which automates experimental phasing for macromolecular structures. Identified selenium atoms were further refined with PHASER^46^. Positions of selenomethionine residues provided key reference points for tracing the polypeptide chain in the electron density map^47^. The CINP-C1ORF109 structure was completed with AutoBuild^48^ in PHENIX^49^ with additional manual model building performed in Coot^50^ based on the experimental electron density map. Final model adjustments involved iterative cycles of model building and refinement with Coot^50^ and PHENIX^49^, ensuring a high-quality structural model. All ΔCINP-C1ORF109 residues were built into the structure, except for the M1 in chains A and C and the truncated N-terminal residues (M1-S17) of CINP. Electron density map and model were depicted using either ChimeraX^51,52^ or Pymol^53^. Data collection, refinement, and validation statistics are shown in **Supplementary Table 5**.

### Cryo-EM sample preparation

250 µL of FLAG-affinity purified 55LCC complex (transiently co-overexpressed in HEK293-6E cells from non-codon optimized expression vectors and purified as described above in Methods section **Purification of 55LCC and SPATA5 variants**) at a concentration of 8-10 mg/mL were incubated with 5 mM ATPγS (Jena Bioscience) for 30 min on ice and loaded onto 10-30% v/v sucrose gradients in buffer A (20 mM HEPES-KOH pH 7.5, 250 mM NaCl, 5 mM MgCl2) and spun for 20 hours at 160,000 rcf in a SW32.1 Ti swing-out rotor (Beckman). Gradients were fractionated using a peristaltic pump (Gilson) coupled to an AKTA Pure 25 system (Cytiva) into a 96-well masterblock (Greiner). Protein peaks were determined by A_280_ absorption and analyzed for their protein content by SDS-PAGE. For stabilization of the complex for cryo-EM, 0.1% v/v glutaraldehyde (Merck) and 200 µM BS3 (bis(sulfosuccinimidyl)suberate) (Thermo Fisher Scientific) crosslinkers were included in the 30% v/v sucrose buffer based on the GraFix^54^ method. Peak fractions from crosslinked gradients were pooled, quenched with 100 mM Tris-HCl pH 8.0 and concentrated to a final concentration of 1 mg/mL. Fresh ATPγS (Jena Bioscience) was added for 30 min on ice at a final concentration of 1 mM before plunge-freezing at the Vitrobot (Mark IV, FEI/Thermo Fisher Scientific) at 95% humidity and 4 °C. For this, 3.5 µL samples were applied to Cu mesh 300 R1.2/1.3 grids (Quantifoil) glow discharged for 45 sec at 15 mA in a Leica Coater ACE 200 glow discharger and grids were plunge-frozen into liquid ethane using a blot force of −8, 5 sec waiting time and 3 sec blotting time before being stored in liquid nitrogen.

### Cryo-EM data collection

Data collection was performed using EPU 2.12.1 (Thermo Fisher Scientific) at a Titan Krios G2 microscope equipped with a Falcon 4i camera and a Selectris energy filter (Thermo Fisher Scientific). The microscope was operated at 300 kV and liquid nitrogen temperature in fast counting mode, an energy filter slid width of 10 eV, and at a nominal magnification of 165,000 with a calibrated pixel size of 0.728 Å. Movies were collected using a total dose of 50 e^-^ at a dose rate of 0.8 e^-^/Å^2^ (50 fractions), spot size 6, and at an applied defocus range of −0.8 to −2.6 μm. For final data processing, data from three different data collections were combined, each of 10,000 movies.

### Cryo-EM data processing and model creation

Data processing was carried out in cryoSPARC 4.2.1^55^. After patch-motion correction and patch CTF correction, about 80% of the pre-processed micrographs were manually selected based on signs of aggregation and ice thickness. Multiple iterative rounds of 2D classification and 3D classification, and heterogeneous refinement were used to clear out conformational heterogeneity within the data set. Shown Fourier Shell Correlation (FSC) curves (**Supplementary Figure S7**) were taken from final homogeneous refinements in cryoSPARC^55^ using the gold standard threshold of 0.143^56^. Selected 3D maps A and B were used to dock AlphaFold2 Multimer^26,27^ generated models of 55L/55LCC complexes manually in ChimeraX^52^ as well as two copies of the biological heterodimer derived from the CINP-C1ORF109 crystal structure. Final models were then created by applying rigid body model docking (using rings of SPATA5/5L1 D1 domains, D2 domains, and N-terminal domains in combination with CINP/C1ORF109 dimers as three separate rigid bodies) and geometry minimization in PHENIX^49^. Cryo-EM maps were depicted using either ChimeraX^52^ or Pymol^53^.

### Polysome profiling and fractionation

HCT116 cell lines were seeded in 500 cm^2^ square cell culture dishes (Corning) at 15 × 10^6^ cells/mL and 10 x 10^6^ cells/mL for the experiment and control, respectively. After cells were grown to ∼40% confluency, the media was exchanged and supplemented with 50 nM PROTAC (dTAG^V^-1) for the experiment dishes or 0.005% DMSO for the control dishes, and the treatment lasted 44 h. Next, cells were incubated with 100 mg/mL cycloheximide for 10 min to arrest translation and harvested by scraping. Pellets were flash-frozen and stored at −80 °C. Cell pellets were subsequently resuspended by pipetting in 1 mL per 0.5 g pellet of ice-cold lysis buffer (20 mM HEPES-KOH pH 7.5, 100 mM NaCl, 5 mM MgCl2, 0.15% NP-40, 100 mg/mL cycloheximide, 1 tablet of cOmplete Inhibitor cocktail EDTA Free (Roche) per 10 mL, RNAse inhibitor (Roche) 2 µL per 10 mL). The cell lysate was cleared by centrifugation at 12,000 rcf for 10 min at 4 °C (FA-24×2 rotor, Eppendorf), and the soluble fraction was collected. The soluble fraction was quantified by nanodrop (A_260_), and 300 µg of RNA was loaded onto a 15% w/v – 45 % w/v sucrose gradient and centrifuged at 180,000 rcf for 3.5 h at 4 °C (SW 32.1 Ti swinging-bucket rotor, Beckman). Gradients were fractionated using a peristaltic pump (Gilson) coupled to an AKTA Pure 25 system (Cytiva) into a 96-well masterblock (Greiner). Protein peaks were analyzed by western blots (iBind system, Thermo Fisher Scientific). The data was plotted using PRISM (v. 9).

### Cell lysis and bottom-up proteomics

Cells were washed twice with ice-cold 1x PBS, followed by on-plate lysis using SDC lysis buffer (2% w/v sodium deoxycholate, 50 mM HEPES, pH 8.0). The resulting lysates were boiled at 95 °C for 15 min, cooled down, and stored at −80 °C until further use. For further use, frozen lysates were thawed, boiled again at 95 °C for 15 min, and sonicated for 2 min or until fully homogenized using a Diagenode Bioruptor Plus for 3×30 sec intervals at high intensity. The sonicated lysates were centrifuged at 16,000 rcf for 15 min, and protein concentration was determined using the BCA Protein Assay Kit (Thermo Fisher Scientific) according to the manufacturer’s instructions. Bottom-up proteomics was performed using different acquisition setups. For data-dependent acquisitions (DDA) based SILAC-based comparisons, equal amounts of protein from the respective comparison groups were pooled for further processing (SILAC-based proteome analysis). Samples for data-independent acquisition (DIA) and pulsed SILAC were processed individually. In all cases, proteins were reduced with 1 mM dithiothreitol (DTT) at 95 °C for 15 minutes. After cooling, samples were alkylated with 55 mM chloroacetamide (CAA) at 30 °C for 1 hour. Next, samples were digested overnight with trypsin at a 1:100 (w/w) enzyme-to-protein ratio at 37 °C with shaking at 800 rpm. Digestion was stopped by adding trifluoroacetic acid (TFA) to a final concentration of 1–1.5% (v/v), followed by centrifugation at 4,800 rcf for 30 minutes to remove sodium deoxycholate. Peptides were purified using Sep-Pak Classic C18 cartridges (Waters) according to the manufacturer’s protocol and eluted with 50% (v/v) acetonitrile (ACN) in MS-grade H₂O. Depending on the experiment, DDA proteomes were subjected to reverse-phase offline fractionation, while pSILAC samples, DIA proteomes, AP-MS, and XL-MS samples were acquired without fractionation. Final eluates were vacuum-concentrated at 60 °C, and peptides were reconstituted in 0.15% TFA at 30 °C with shaking at 1,500 rpm for 15 minutes. Peptide concentrations were determined using a NanoDrop before MS acquisition. An overview of all MS-Experiments and resulting data as well as experimental details and used cell lines, is provided in **Supplementary Table 6**. An overview of all MS-Settings is provided in **Supplementary Table 7**.

### Affinity purification mass spectrometry

Cells expressing affinity-tagged (GFP / mCherry / 3xFLAG / HaloTag) POI were cultured in ‘heavy’ or ‘medium’ isotope-containing SILAC media. Cells expressing free GFP were labeled with ‘light’ SILAC isotopes and used for control pull-downs to estimate background binding proteins. Cells were washed three times with ice-cold 1x PBS, scraped off, pelleted at 1,000 rcf for 1 minute at 4 °C, and lysed in RIPA buffer (50 mM Tris-HCl pH 7.5, 150 mM NaCl, 1% NP40, 0.1% sodium deoxycholate, 1 mM EDTA pH 7.5, 5 mM β-glycerophosphate, 5 mM sodium fluoride, 1 mM sodium orthovanadate, and cOmplete Inhibitor cocktail EDTA Free (Roche) while rotating at 4 °C for 30 min. Next, lysates were sonicated three times for 30 sec at high intensity at 4 °C using a Diagenode Bioruptor Plus, followed by centrifugation at 16,000 rcf for 10 min at 4 °C. Supernatants were transferred to new tubes, and protein concentrations were measured using the Pierce BCA assay on a Tecan Spark system. Protein concentrations were adjusted to a maximum of 2 mg/mL, and 2 mg of protein was used for each affinity purification (AP) condition. Next, prewashed beads specific for the fused TAGs were added to the lysates (15 µL GFP-trap Magnetic Agarose, Chromotek/Proteintech (gtma-200); 15 µL RFP-trap Magnetic Agarose, Chromotek/Proteintech (rtma-200); 20 µL anti-FLAG M2 Magnetic Beads, Sigma, (M8823-1ML); and 15 µL HaloTrap-beads (otma-20)), and the affinity enrichment was carried out for 90 min at 4 °C with rotation at 10 rpm. Beads were washed once with 1 mL of RIPA buffer, and control and bait affinity-enriched samples were combined at this step. After two additional washes with AP-MS wash buffer (10 mM Tris-HCl pH 7.5; 150 mM NaCl; 0.5 mM EDTA), proteins were digested on-bead using 50 µL of elution buffer 1 (50 mM Tris-HCl pH 7.5, 2 M urea, 5 µg/mL trypsin, 1 mM DTT) for 30 min at 30 °C with shaking at 800 rpm. The supernatants were collected together with two additional bead washes using 100 µL of elution buffer 2 (50 mM Tris-HCl pH 7.5, 2 M urea, 5.5 mM CAA). The eluted protein mixtures were incubated overnight at 37 °C with shaking at 1,000 rpm. The next day, peptides were purified using stage tips with three C18 discs, vacuum-concentrated at 60 °C, and finally dissolved in 0.15% TFA at 30 °C with shaking at 1,500 rpm for 15 min. Peptide concentrations were measured using a NanoDrop One (Abs1 = 1 mg/mL), and 200-300 ng of peptides per sample were used for subsequent MS analysis.

### In gel sample preparation

After affinity-enrichment, the beads were washed three times with ice-cold RIPA lysis buffer. Proteins bound to the beads were then eluted by heating the beads at 95 °C for 15 min in 50 µL of 4x NuPAGE LDS sample buffer (Invitrogen) containing 1mM DTT. Afterward, 1 µL of 550 mM chloroacetamide (CAA) was added, and the samples were incubated at room temperature (RT) for 30 min. The eluted proteins were then subjected to in-gel digestion for MS analysis. First, the proteins were separated on a NuPAGE Novex Bis-Tris 4%-12% gel (Invitrogen). The gel was stained with Novex colloidal blue stain (Invitrogen) and destained with water. Each lane was cut into four slices, which were further destained using 25 mM ammonium bicarbonate buffer containing 50% ethanol. The gel pieces were then dehydrated with 100% ethanol before being digested in-gel with trypsin (Sigma-Aldrich) at 37 °C for 16 h. Following digestion, the gel pieces were treated with trifluoroacetic acid (TFA) to stop the reaction, and the resulting peptides were eluted using increasing concentrations of acetonitrile. The resulting peptides were desalted using C18 StageTips. For elution from the StageTips, 50 µL of a 60% acetonitrile and 0.1% TFA solution was used. The peptides were then vacuum-dried at 60 °C. Dried peptides were resuspended in 0.15% TFA buffer at 30 °C and shaken at 1,500 rpm for 15 min. The resulting peptide concentration was estimated using a NanoDrop and 300 ng of the sample was used for subsequent MS analysis.

### Pulsed SILAC (pSILAC) for protein synthesis capability

pSILAC was performed to quantify the amount of newly synthesized proteins, reflecting the rate of protein synthesis^23^. Before the assay, cells were cultured in ‘light’ SILAC media. Degron-tagged cell lines were either pre-treated with DMSO or pre-depleted (for 12, 24, 48 hours before the start of pulse-SILAC labeling) of 55LCC proteins using 50 nM dTAG^V^1. Cells were pulse-SILAC labeled by switching to ‘heavy’ SILAC media for 24 h duration, while maintaining their respective treatments. Residuals of the original media were removed by washing the cells three times with ‘heavy’ SILAC media. After 24 h, the assay was stopped, and the cells were processed as previously described via the SDC lysis and processing protocol. In this setup, the detected protein IDs in heavy SILAC represent newly synthesized proteins during the pulse, thereby reflecting the protein synthesis rate/capacity of the tested cells. DMSO pre-treated cells served as reference controls, and the detected protein IDs were compared relative to each other.

### Immunocytochemistry image analysis

After unbiased automated image acquisition, quantitative image analysis was performed in an automated manner using the PerkinElmer Harmony v. 5.2 software, with data processed through experiment-specific analysis pipelines. The resulting output data were analyzed in R. Representative micrographs were pseudo-colored and displayed in the manuscript without further modification other than cropping, while image export was done with the same setup as used for the controls. For image analysis, Flatfield and Brightfield corrections were applied to image acquisitions across all channels, planes, and fields using the advanced options. For analyses of Z-stack acquisitions, maximum intensity projections were generated for subsequent analysis. Background correction was applied at fixed parameters before further processing. Hoechst-stained Nuclei were identified in an automated manner. Cytoplasm was detected automatically as well, by the stained signal area with respect to detected nuclei, both together enabling discrimination of nuclear vs. cytoplasmic cell area, using the built-in detection methods. Consistent detection mode, threshold, area, splitting coefficient, individual threshold, and contrast parameters were applied within each experiment and quantification condition.

For detecting changes in subcellular distributions of ribosome biogenesis factors after perturbation of 55LCC constituents, antibody signals of the respective proteins were analyzed in nuclear and cytoplasmic areas, based on the above-described approach, normalized to the size of the analyzed cell area. Data of the detected intensities were extracted, and the ratio was compared between nuclear and cytoplasmic areas. For all analyses, median intensities were used from the area analyzed. Signal intensities in the DMSO control experiment were used to determine baseline levels and to normalize and log2-transform the data. For display of the micrographs, optimal levels were applied in the control channel, and these settings were fixed for all exports.

### MS data processing

For DDA experiments, raw data were processed using MaxQuant^57^ (v. 2.0.1.0), with searches conducted against the UniProt databases for *Homo sapiens*. The false discovery rate (FDR) for both peptides and proteins was maintained at 1%. Cysteine carbamidomethylation was applied as a fixed modification, while methionine oxidation and N-terminal acetylation were set as variable modifications. Re-quantification was enabled, and the number of allowed missed cleavages was set to two. All other parameters were left at their default settings.

For the analysis of DIA raw data, we utilized DIA-NN^58^ (v. 1.8.1) to predict a spectral library with the following parameters: a maximum of 2 missed cleavages, up to 2 variable modifications, peptide lengths between 7 and 30 residues, precursor charge states ranging from 1 to 4, and precursor m/z and fragment ion m/z ranges of 300-1000 and 100-2000, respectively. The following modifications were accounted for: N-terminal methionine excision, oxidation of methionine, N-terminal acetylation, and carbamidomethylation of cysteine. DIA raw files were converted to mzXML format using MSConvert^59^ v. 3.0.24007-5bacee6 (ProteoWizard) and analyzed using the predicted library in DIA-NN^58^ (v. 1.8.1) with the following options enabled: isotopologue utilization, match between runs (MBR), heuristic protein inference, and exclusion of shared spectra. Mass accuracy and MS1 accuracy were set to 15 ppm across all runs. Additional settings included: protein inference set to genes, neural network classifier in single-pass mode, quantification strategy set to Robust LC (high accuracy), cross-run normalization based on RT-dependence, and library generation utilizing smart profiling. Speed and RAM usage were configured for optimal performance.

### Processing and analysis of DDA data

In R^60^ (v. 4.3.1), the ProteinGroups table from the MaxQuant^57^ output was filtered to exclude entries originating from the reverse decoy database, contaminants, and proteins identified solely by a modification site. Relative quantification of protein fold changes was determined using SILAC ratios extracted from the ProteinGroups table. Statistical significance was assessed for proteins identified in at least three biological replicates. Differential protein abundance was estimated by a two-sided test using the limma package^61^. P-values below ε_machine_ are automatically set to 0 by limma, so all p-values of 0 were replaced by ε_machine_ (2.220446e^-16^). The Benjamini-Hochberg method was applied for multiple comparison correction. Scatter plots were generated by plotting the negative log10-transformed p-values against the log2-transformed fold changes between the perturbation and control. Proteins with a fold change of ≥1 and a p-value ≤0.05 were considered significantly enriched.

### Data processing of DIA label-free quantification

The MSStats R package^60^ was employed to estimate protein abundances from the report.tsv output file generated by DIA-NN^58^, using the dataProcess function using MSstats. For differential protein abundance analysis, linear mixed-effect models were fitted via the groupComparison function in MSStats^60^, applied to the processed data. Bar plots represent the mean, with error bars indicating the standard error of the mean.

### Processing and network display of SILAC-AP-MS data

To map endogenous interactomes from all SILAC-AP-MS experiments, we generated a unified dataset of all identified proteins across bait conditions (**Supplementary Table 8**). Gene names were standardized to one representative per protein and annotated using DepMap-derived essentiality scores (**Supplementary Table 3**) and a curated list of known ribosomal proteins (**Supplementary Table 4**). To reduce background from abundant or nonspecific binders, we excluded histones and mitochondrial ribosomal proteins using keyword filters (“histone”; “mitochondrial” + “ribosomal”). These proteins are frequently detected in pulldown and proximity-labeling assays due to high abundance or charge properties, but do not reflect specific interactions. The curated dataset (**Supplementary Table 9**) was used for downstream visualization and analysis. For display, we focused on significantly enriched interactors (mean log₂ fold change ≥ 1) classified as essential in at least 50% of DepMap^14^ cell lines (Essentiality Score ≥ 0.5). To highlight potentially novel interactions and reduce network complexity, we removed known interactions annotated in STRING^15^ with confidence scores < 0.5. For proteins found in multiple bait conditions, we retained only the instance with the highest mean log2 fold change. Interactors without prior links to ribosomes were prioritized for follow-up.

### Data processing and analysis of Non-Mass Spectrometry data

All experiments were performed at least three times, and unless specified, no outliers were removed from any of the experiments. The resulting output data were plotted and statistically analyzed using either R^60^ (v. 4.3.1) or PRISM (v. 9). The names of statistical tests used for individual comparisons are indicated in the relevant figure legends. For multiple testing correction, Benjamini-Hochberg or Šidák correction was applied.

## Acknowledgments

The Novo Nordisk Foundation Center for Protein Research is financially supported by the Novo Nordisk Foundation (NNF14CC0001). The Novo Nordisk Foundation Center for Stem Cell Biology is supported by the Novo Nordisk Foundation (NNF15OC0017774). C.C. was supported by a grant from the Independent Research Fund Denmark (10.46540/2034-00429B). G.M. was supported by the ERC-AdG 101096548 (INTETOOLS), NNF0024386, NNF17SA0030214, and NNF18OC0055061 grants. G.M. is a member of the Integrative Structural Biology Cluster (ISBUC) at the University of Copenhagen. We thank the CPR Mass Spectrometry Platform and the CPR Imaging Platform for their assistance. We thank the Danish Cryo-EM National Facility at the University of Copenhagen for support during data collection. We thank the members of our laboratories for their helpful discussions.

## Author contributions

Conceptualization, C.C., J.W. and G.M.; methodology, M.W., J.W., M.M., and G.M. (in detail: cell culture – M.W., J.W., T.W., M.M. and E.B.N.; cloning and expression of proteins in mammalian and bacterial cells – M.W., M.M., E.B.N., R.G.M., N.Z.; protein purification – M.W., M.M. and R.G.M.; crystallization, X-ray data collection – M.W. and M.M.; X-ray data processing, atomic model building and refinement – M.W., M.M. and G.M.; cryoEM sample preparation, data collection, data processing and model creation – M.W.; polysome profiling and fractionation – M.W., M.M. and N.Z.; generation of CRISPR edited cell lines, immunofluorescence cytochemistry imaging, high-throughput microscopy, bright-field image-based growth assessment, proteomics, SILAC, pSILAC, MS, AP-MS – J.W., T.W.; data analysis – M.W., J.W., M.M., T.L., T.N., and G.M.; resources, J.W., C.C. and G.M.; writing – original draft, M.W., and M.M.; writing – review & editing, M.W., J.W., M.M. and G.M.; visualization, M.W., J.W. and M.M.; supervision, project administration, and funding acquisition, C.C and G.M.

## Data availability

The mass spectrometry data have been deposited to the ProteomeXchange Consortium via the PRIDE^62^ partner repository as separate datasets with the following identifiers: AP-MS (PXD065526), pSILAC (PXD064721), DDA Proteomes (PXD064896), DIA Proteomes (PXD064619). Crystal structure of CINP-C1ORF109 complex is available in Protein Data Bank under the 9GWJ accession code. The 55LCC complex map is available in EMDB under the EMD-55042 accession code.

## Experimental Design

All experiments were performed in replicates. No aspect of the study was done blinded. Sample size was not predetermined, and no outliers were excluded.

## Supplementary figures

**Figure S1.**
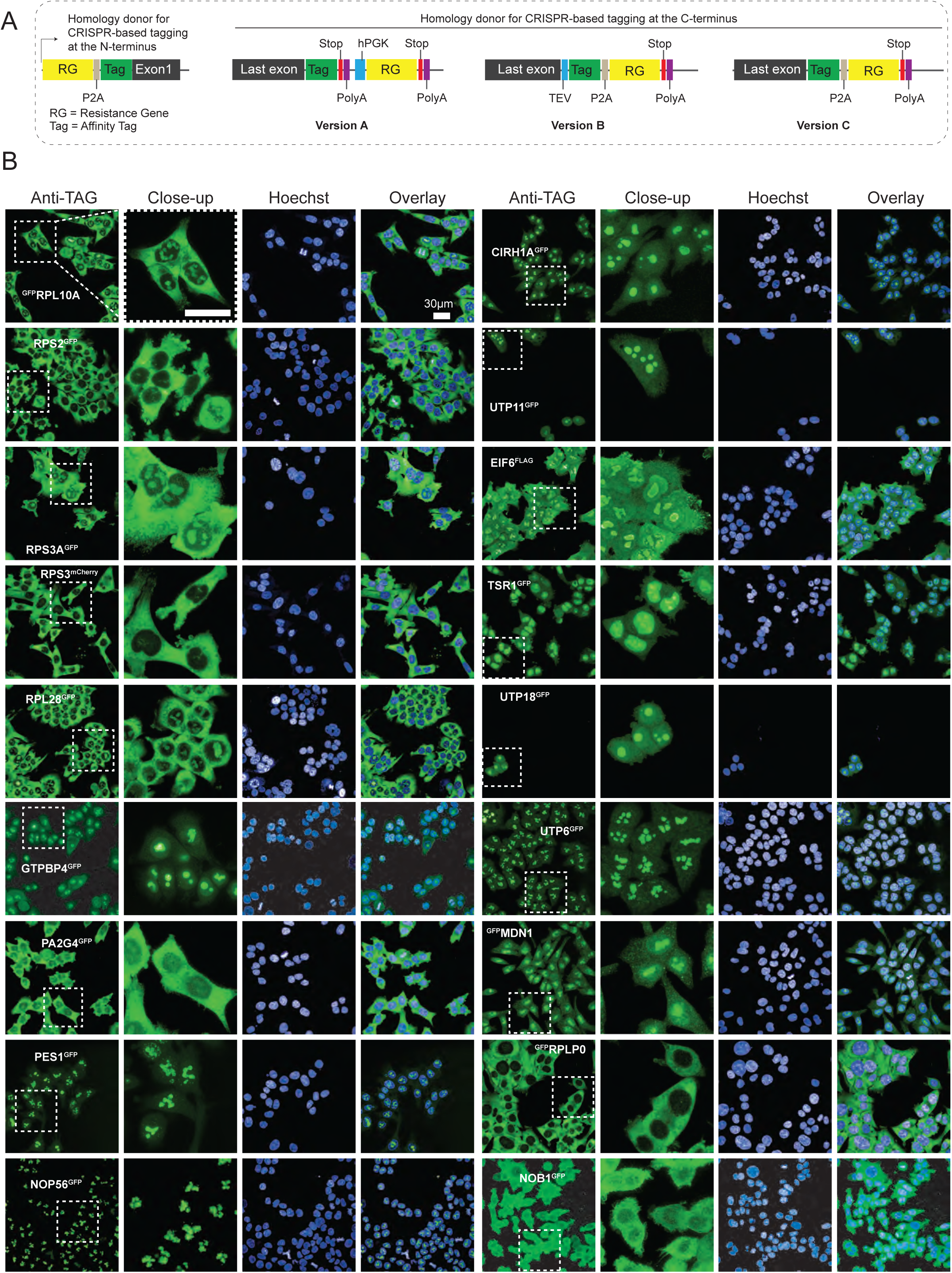
Design for endogenous tagging and confirmation of tagged protein expression. (**A**) Schematic display of editing cassettes used for endogenous CRISPR editing of cell lines used for the systematic interactome mapping. (**B**) Microscopy images showing subcellular localization of the affinity-tagged bait proteins. UTP18^GFP^, UTP6^GFP^, ^GFP^MDN1, and NOB1^GFP^ were expressed but affinity enrichment did not yield expected interactors and hence were not used for further analyses.

**Figure S2.**
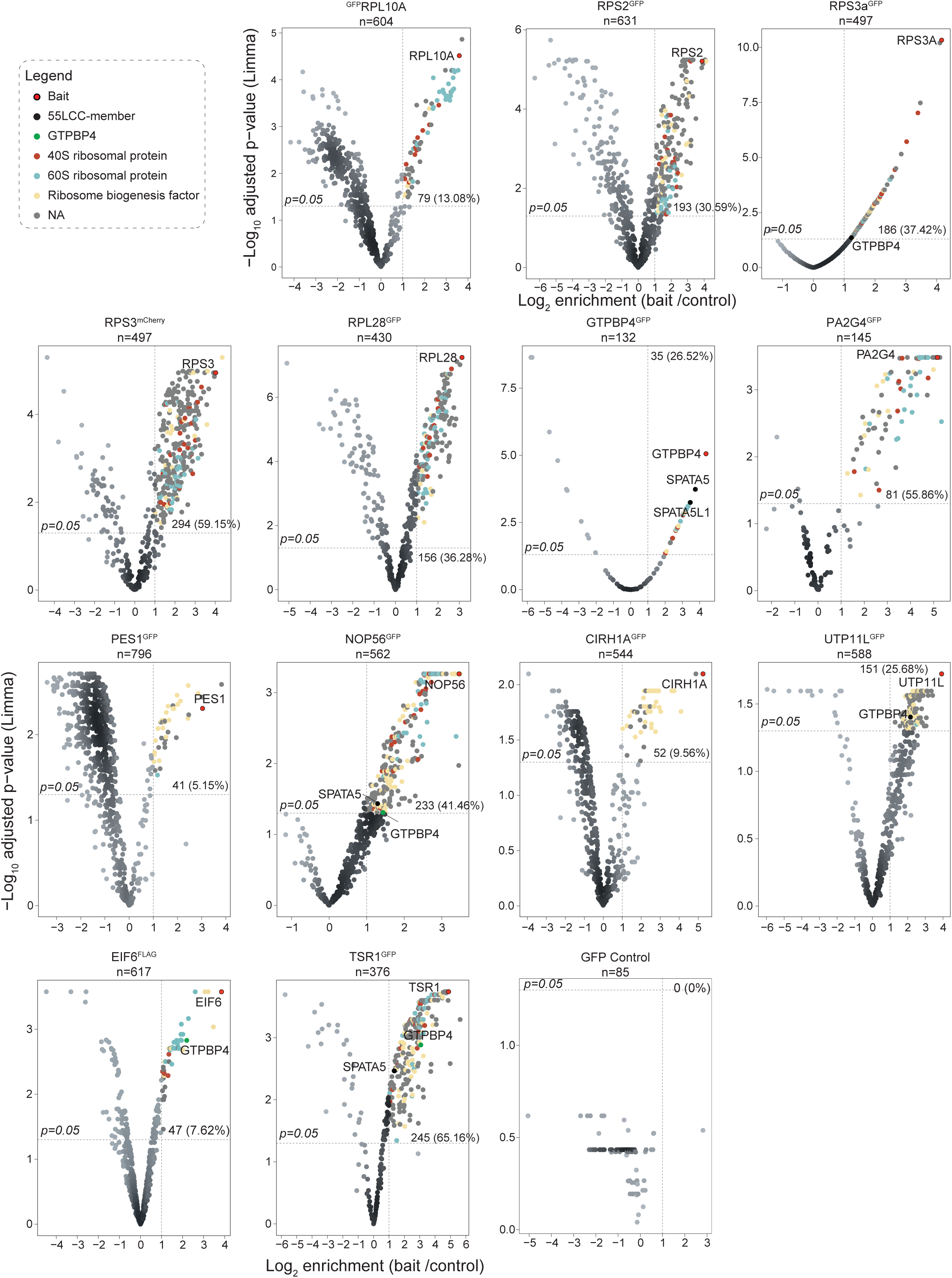
Mapping interactomes of affinity-tagged ribosome-associated factors and ribosomal proteins. Volcano plots displaying LIMMA-derived significance (–log₁₀ adjusted p-values, y-axis) versus fold-enrichment (log₂ fold-change, x-axis) of identified proteins (*n*≥3). Each plot indicates the bait protein, total number of detected proteins (top), and the number and percentage of significantly enriched interactors. A horizontal line marks the p = 0.05 significance threshold. Shown are interactomes from bait cell lines selected for downstream network analysis, alongside the interactome of free GFP-expressing cells used as a background control. The legend highlights color-coded categories of mapped interactor groups, bait proteins, or specific interactors.

**Figure S3.**
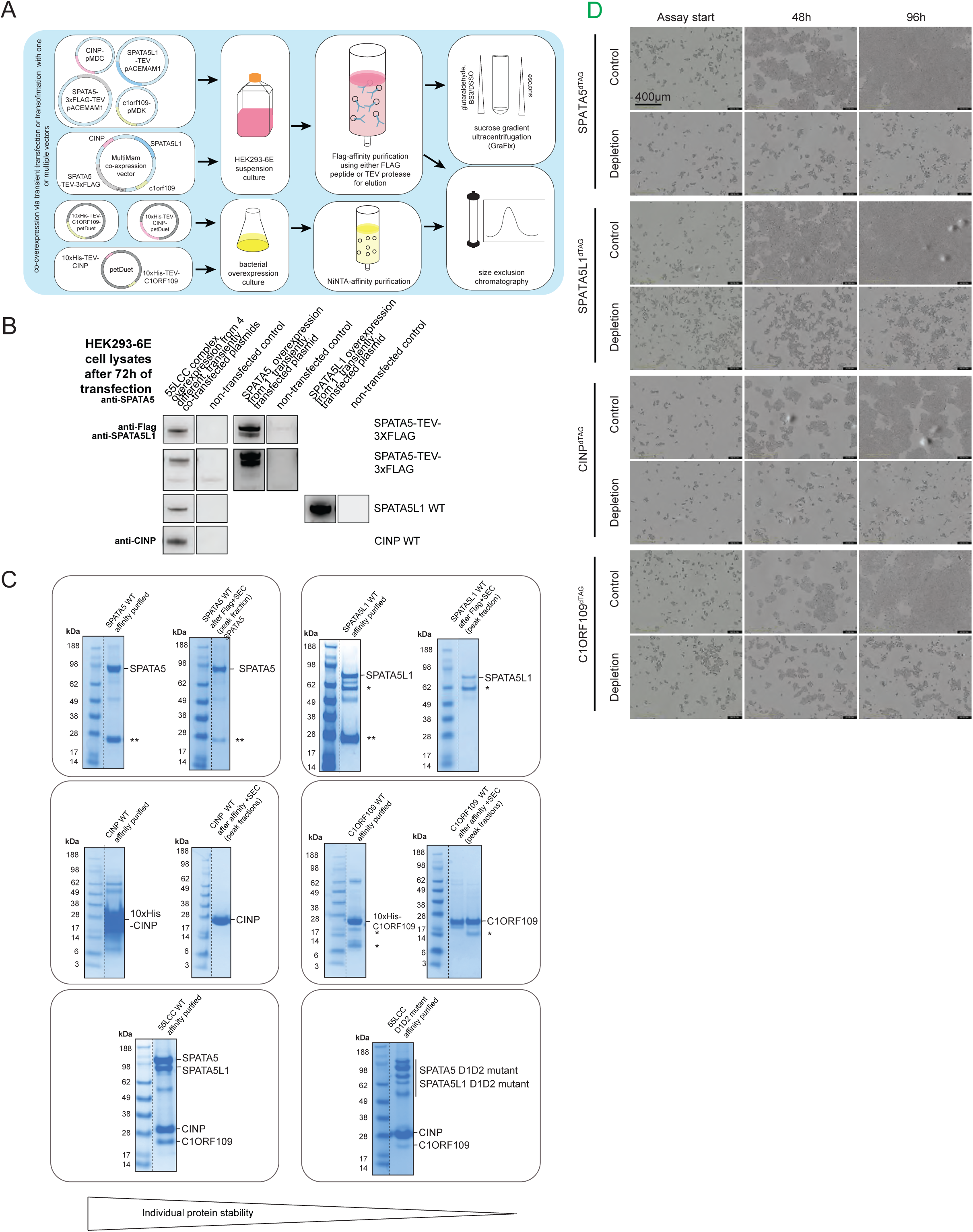
AP-MS data processing, screen validation, and functional validation of endogenously degron-tagged 55LCC proteins. (**A**) Outline of the production strategies used for expressing and purifying recombinant human SPATA5-SPATA5L1-CINP-C1ORF109 and CINP-C1ORF109 complexes, as well as individual CINP, C1ORF109, SPATA5 and SPATA5L1 proteins. (**B**) Western blots demonstrating co-overexpression of human 55LCC complex in HEK293-6E cells, as well as human SPATA5 and human SPATA5L1 after 72h of transient transfection. (**C**) Coomassie-stained SDS-PAGEs of recombinantly expressed and affinity and SEC-purified SPATA5, SPATA5L1, as well as CINP and C1ORF109 proteins. Under the same expression and purification conditions, SPATA5L1 and C1ORF109 are more prone to protein degradation than SPATA5 and CINP, resulting in lower overall protein yields. Besides, SDS-PAGE analysis of recombinantly expressed and purified 55LCC wild-type complexes shows that SPATA5L1, as well as C1ORF109, are stabilized when incorporated in the 55LCC complex. (**D**) Representative brightfield images used for monitoring cell growth during dTAG^V^1 exposure. Solvent control and dTAG^V^1-treated cells of the same line are shown side-by-side for direct comparison. Statistical significance intervals indicated as: * p<0.05, ** p<0.01, *** p<0.001, **** p<0.0001.

**Figure S4.**
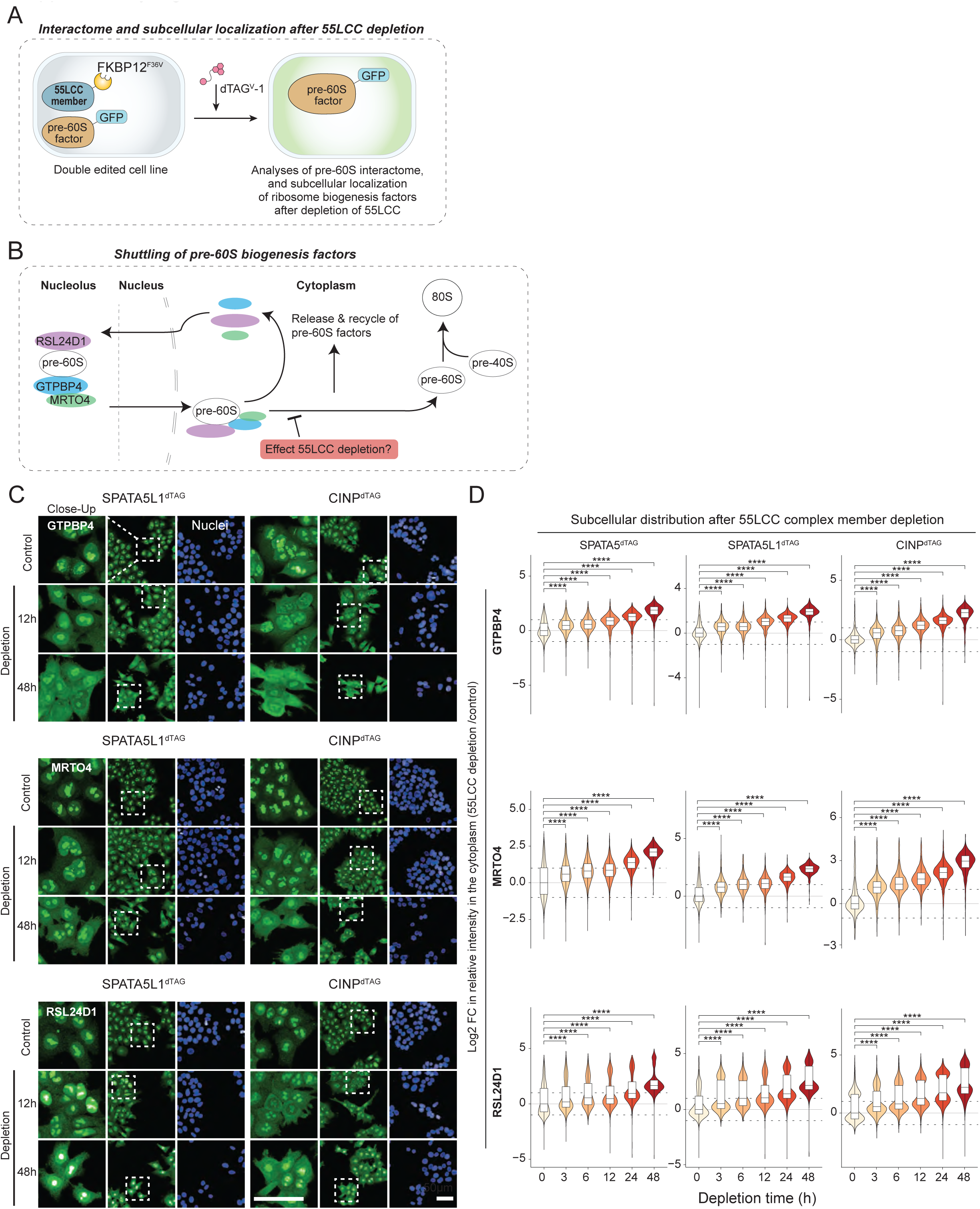
55LCC depletion impairs the release of pre-60S-associated ribosome biogenesis factors in the cytoplasm. (**A**) Schematic presentation of the strategy used for mapping pre-60S interactome after the depletion of 55LCC complex members. First, endogenous 55LCC constituents were fused with FKBP126^F36V^ (dTAG), and in this background, pre-60S-binding factors (GTPBP4, RPL28, and RPL10A) were fused with GFP in their endogenous loci. Interactomes of pre-60S-binding factors were analyzed with or without depletion of individual 55LCC constituents (see **Figure 4A-B**). (**B**) Schematic representation of the release of the indicated pre-60S biogenesis factors in the cytoplasm and their recycling in the nucleolus. (**C**) Micrographs of GTPBP4, MRTO4, and RSL24D1 immunocytochemistry in untreated conditions and after the depletion (12h, 48h) of SPATA5L1^dTAG^ or CINP^dTAG^. (**D**) Quantification of fluorescence intensities of the indicated pre-60S biogenesis factors in specified subcellular compartments at the indicated timepoints after the depletion of SPATA5^dTAG^, SPATA5L1^dTAG^ or CINP^dTAG^ (*n*=3). Violin plots display the distribution of log₂-transformed intensity ratios between cytoplasm and nucleus across treatment time points. Boxplots are overlaid within each violin and indicate the median (horizontal bar) and the interquartile range (IQR; box hinges at the 25th and 75th percentiles). Whiskers extend to 1.5× the IQR; outliers beyond this range are not shown. Statistical comparisons to the untreated control (0 h) were performed using a two-sided Wilcoxon test with Benjamini–Hochberg correction for multiple testing **** p<0.0001. Dotted lines at ±1 log2 fold change.

**Figure S5.**
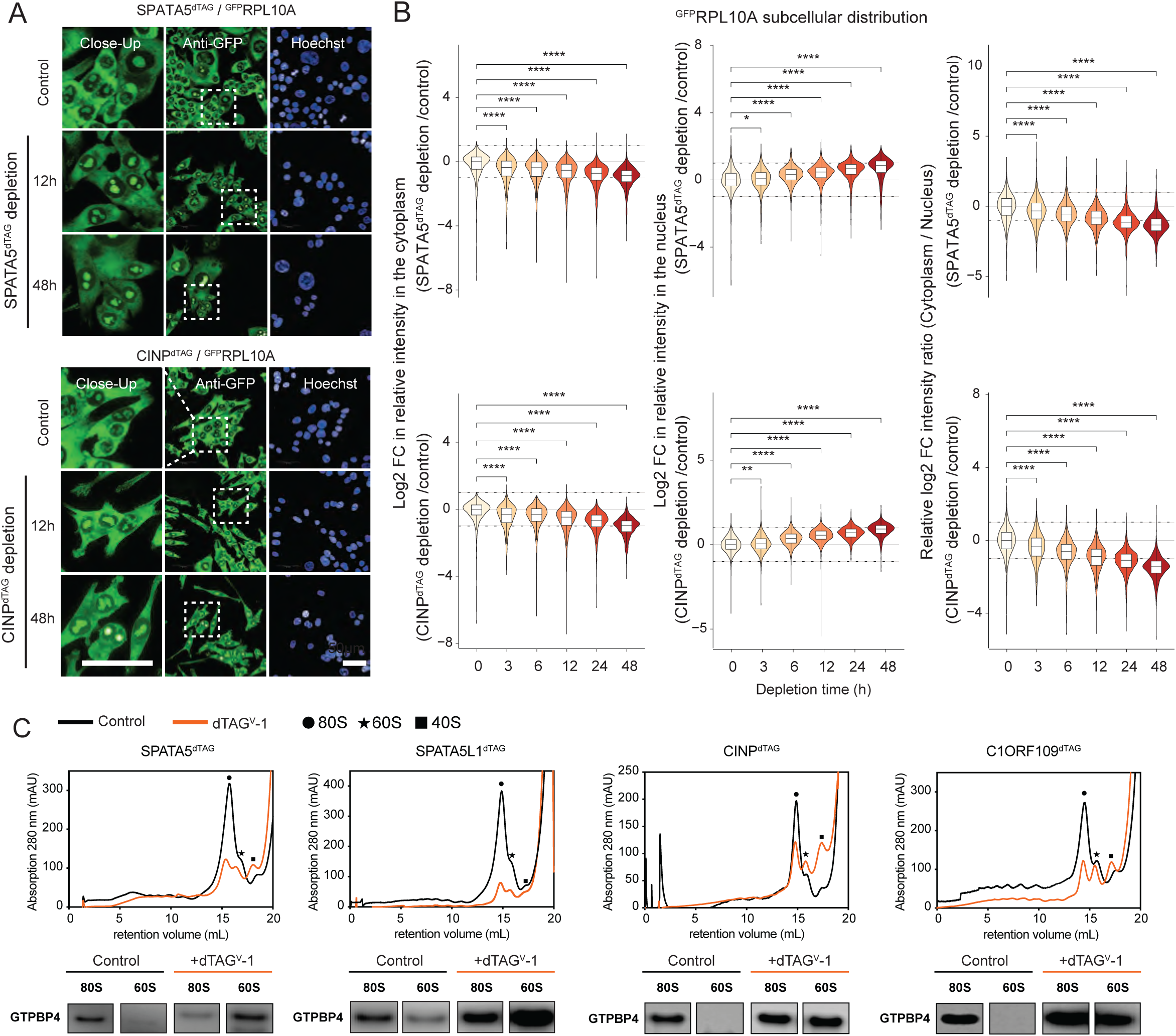
55LCC mutant cells display ribosome assembly defects and altered RLP10A sub-cellular localization after the depletion of SPATA5^dTAG^ or CINP^dTAG^. (**A**) Micrographs of ^GFP^RLP10A in untreated condition and after the depletion (12h, 48h) of SPATA5^dTAG^ or CINP^dTAG^. (**B**) Quantification of fluorescence intensities of ^GFP^RLP10A in specified subcellular compartments at the indicated timepoints after the depletion of SPATA5^dTAG^ or CINP^dTAG^ (*n*=3). Violin plots display the distribution of log₂-transformed intensity ratios between cytoplasm and nucleus across treatment time points. Boxplots are overlaid within each violin and indicate the median (horizontal bar) and the interquartile range (IQR; box hinges at the 25th and 75th percentiles). Whiskers extend to 1.5× the IQR; outliers beyond this range are not shown. Statistical comparisons to the untreated control (0 h) were performed using a two-sided Wilcoxon test with Benjamini–Hochberg correction for multiple testing **** p<0.0001. Dotted lines at ±1 log2 fold change. (**C**) Polysome profiling and analysis of cell lines with depleted wild-type 55LCC components. For each component, a representative polysome profile from the control culture is shown in black and compared to a polysome profile from cultures subjected to 44 hours of protein depletion, shown in blue (top panels). Fractions were analyzed by SDS-PAGE and Western blot using an antibody against GTPBP4 (bottom panels).

**Figure S6.**
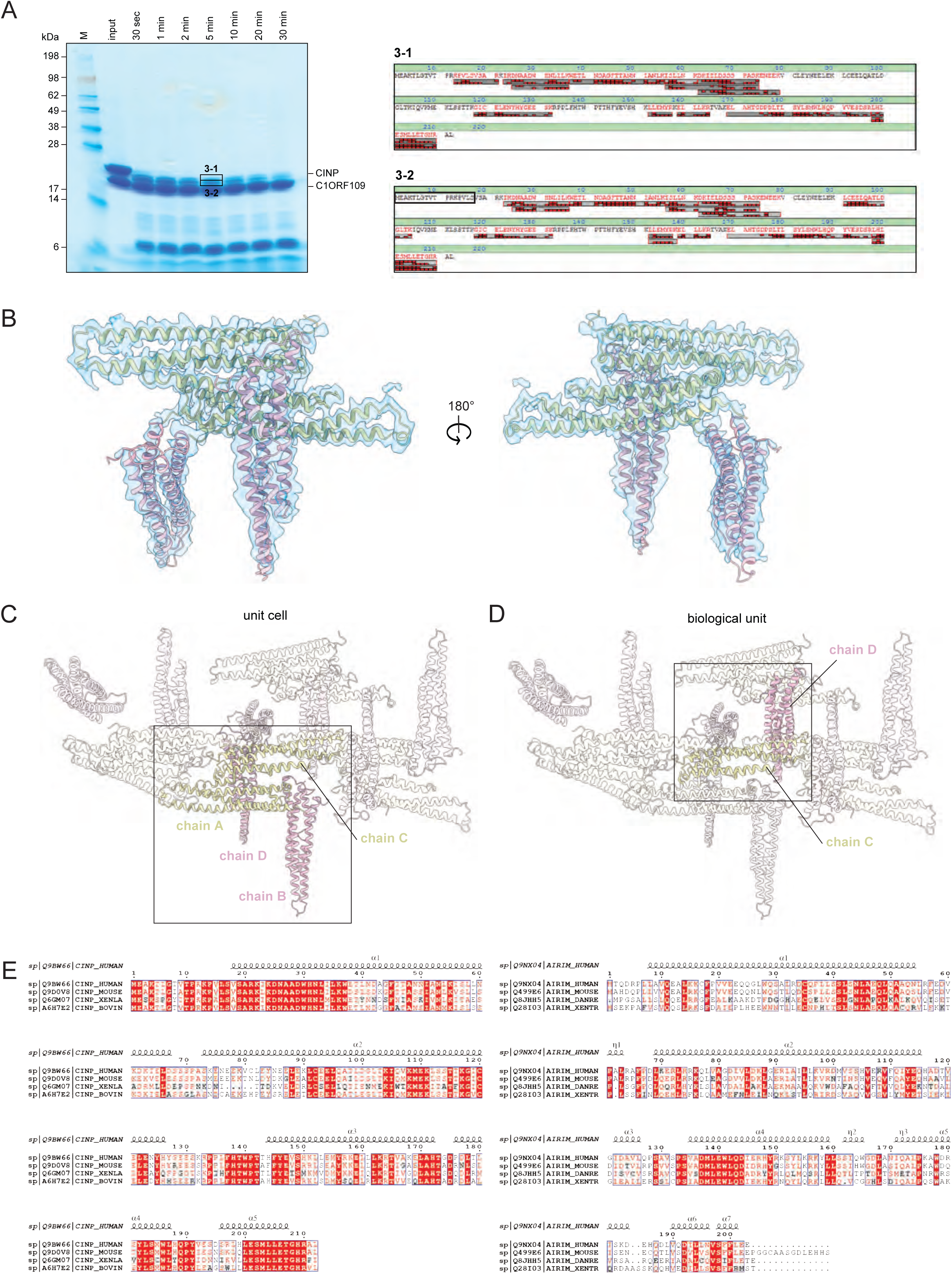
Limited proteolysis mapping and structural fitting of CINP-C1ORF109 complex. (**A**) Left panel, limited proteolysis experiment indicating significant degradation of the N-terminal tail of CINP. Coomassie-stained SDS-PAGE analysis of full-length CINP-C1ORF109 complex after treatment with trypsin. CINP degrades into multiple bands, while C1ORF109 appears stable with a single band. Right panel, the N-terminal peptide cleaved by trypsin is marked with a black box on the mass spectrometry report. (**B**) The CINP-C1ORF109 complex model fitted into the electron density map. 2Fo–Fc map is presented as a blue transparent surface, contoured at 1.0 σ. (**C**) Crystal packing of chains A, C (yellow), and chains B, D (pink). The unit cell is shown as a solid cartoon and marked with a black box; symmetry-related molecules are represented as a transparent cartoon. (**D**) The biological unit of the CINP-C1ORF109 complex is formed between chain D of the unit cell and chain C of a symmetry-related molecule. The biological unit is depicted as a solid cartoon and marked with a black box. (**E**) Multiple sequence alignment of CINP (left panel) and C1ORF109 (right panel) wild-type protein sequences created by T-coffee multiple sequence alignment server^63^ and ESPript^64^ v. 3.0.

**Figure S7.**
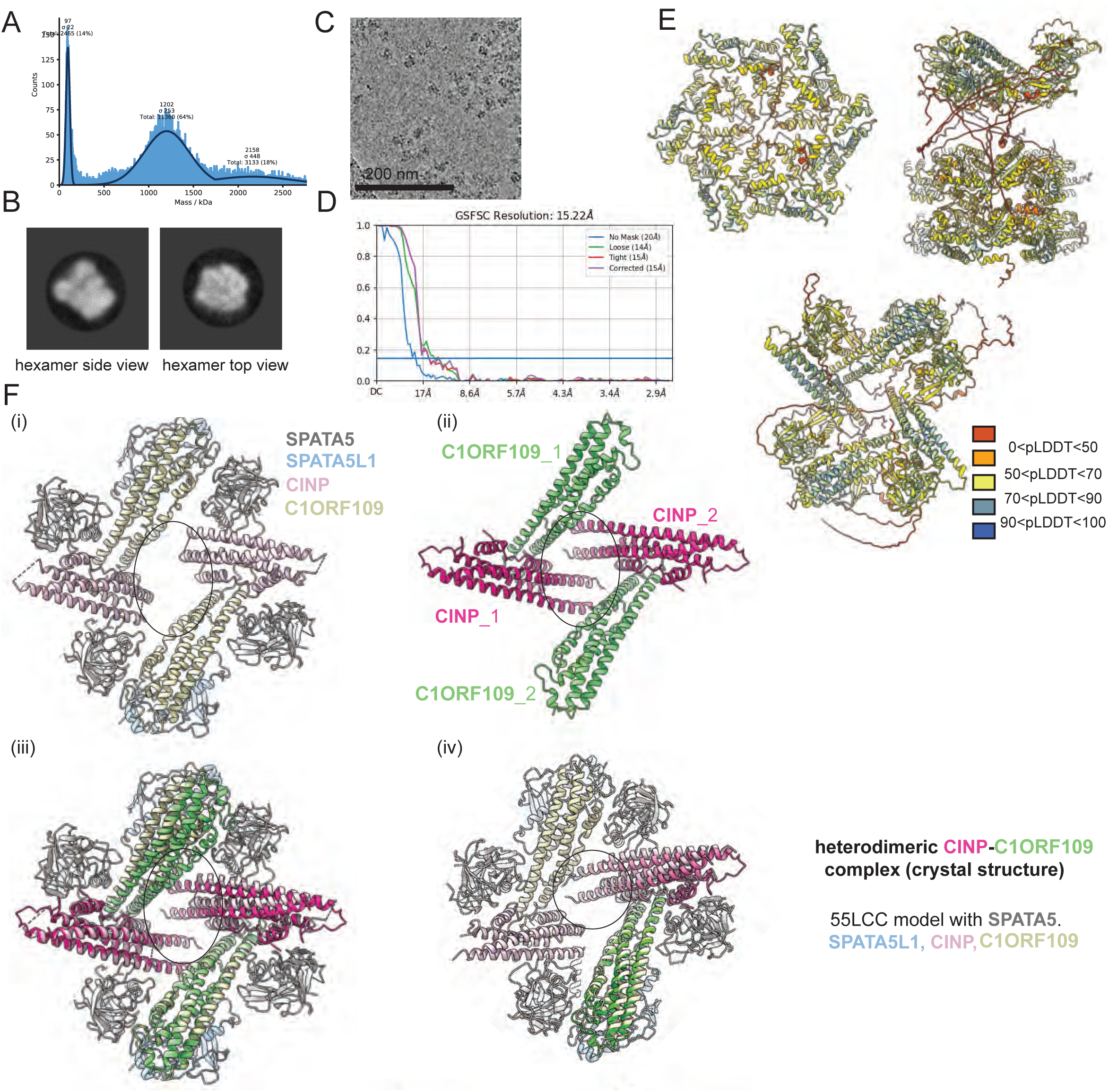
Biophysical characterization and structural modeling of the 55LCC complex. (**A**) Particle size distribution of FLAG affinity-purified CINP/C1ORF109/SPATA5/SPATA5L1 complexes in the absence of nucleotide determined by mass photometry. (**B**) 2D class averages determined by negative stain transmission electron microscopy and data processing in cryoSPARC^55^ represent characteristic side and top views of the 55LCC complex isolated from a GraFix^54^ gradient peak fraction, allowing identification of the heterohexameric architecture and stoichiometry of the ATPase complex. (**C**) Example micrograph of the 55LCC complex isolated from GraFix^54^ gradient peak fraction c determined by cryo-EM, showing asymmetric particle distribution in the grid holes. (**D**) FSC curve and overall resolution of the 55LCC cryo-EM map B used for 55LCC structural modeling. (**E**) AlphaFold 2 Multimer model^26,27^ of a 55LCC complex colored by pLDDT value. Despite the on-average low per-residue confidence, the predicted model is in good agreement with the experimentally observed complex architecture. (**F**) Structural comparison of the CINP-C1ORF109 heterodimers within the 55LCC complex model (i) with the CINP-C1ORF109 heterodimer determined by X-ray crystallography (ii). Two copies (iii) or one copy (iv) of the heterodimer from the crystal structures were aligned and superimposed with ChimeraX Matchmaker^51,52^.

**Figure S8.**
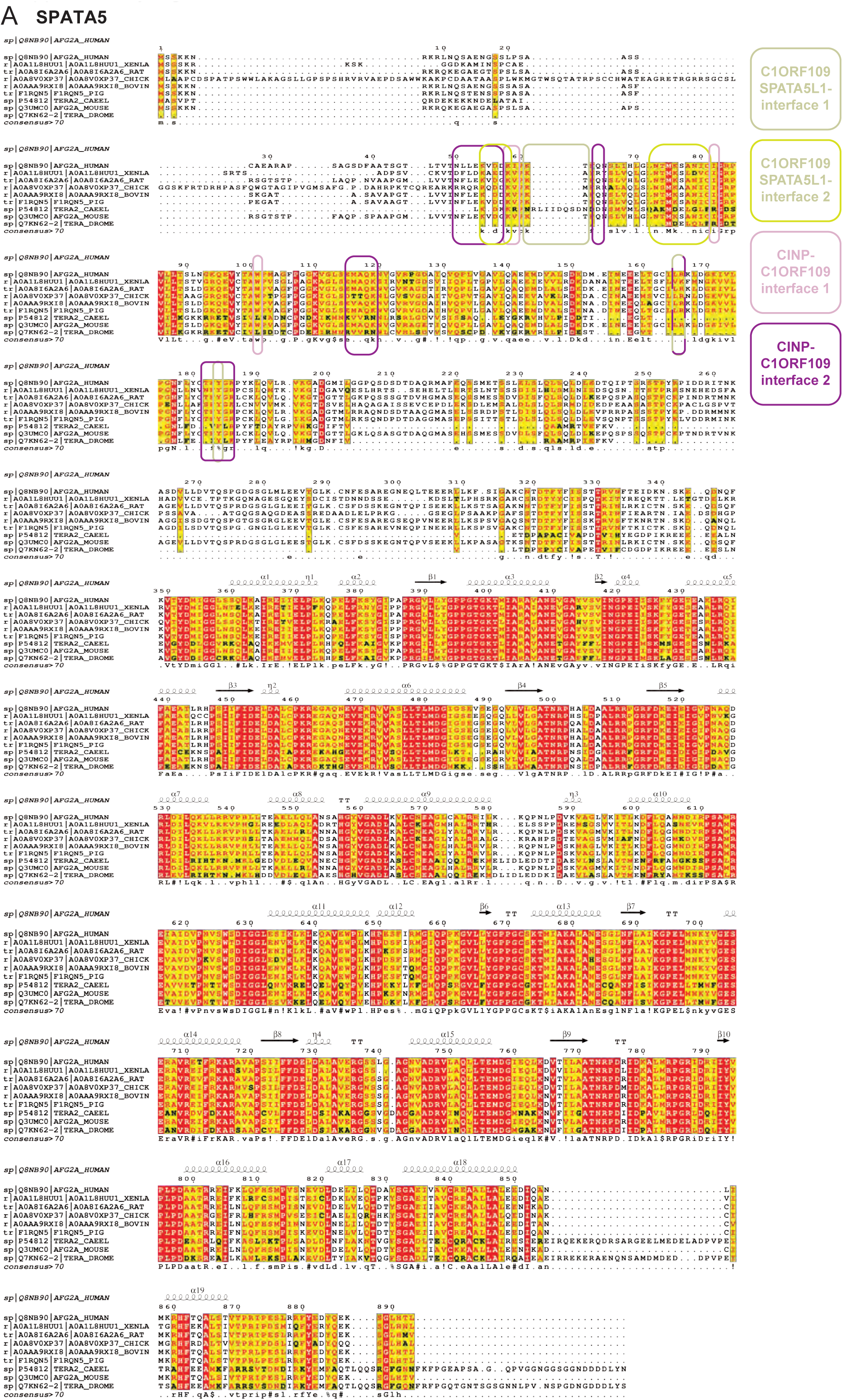
Multiple sequence alignment of SPATA5 wild-type protein sequences. Alignment created by T-coffee multiple sequence alignment server^63^ and ESPript v. 3.095^64^. Interface areas with cofactor CINP-C1ORF109, as determined by cryo-EM and structural modeling, are indicated.

**Figure S9.**
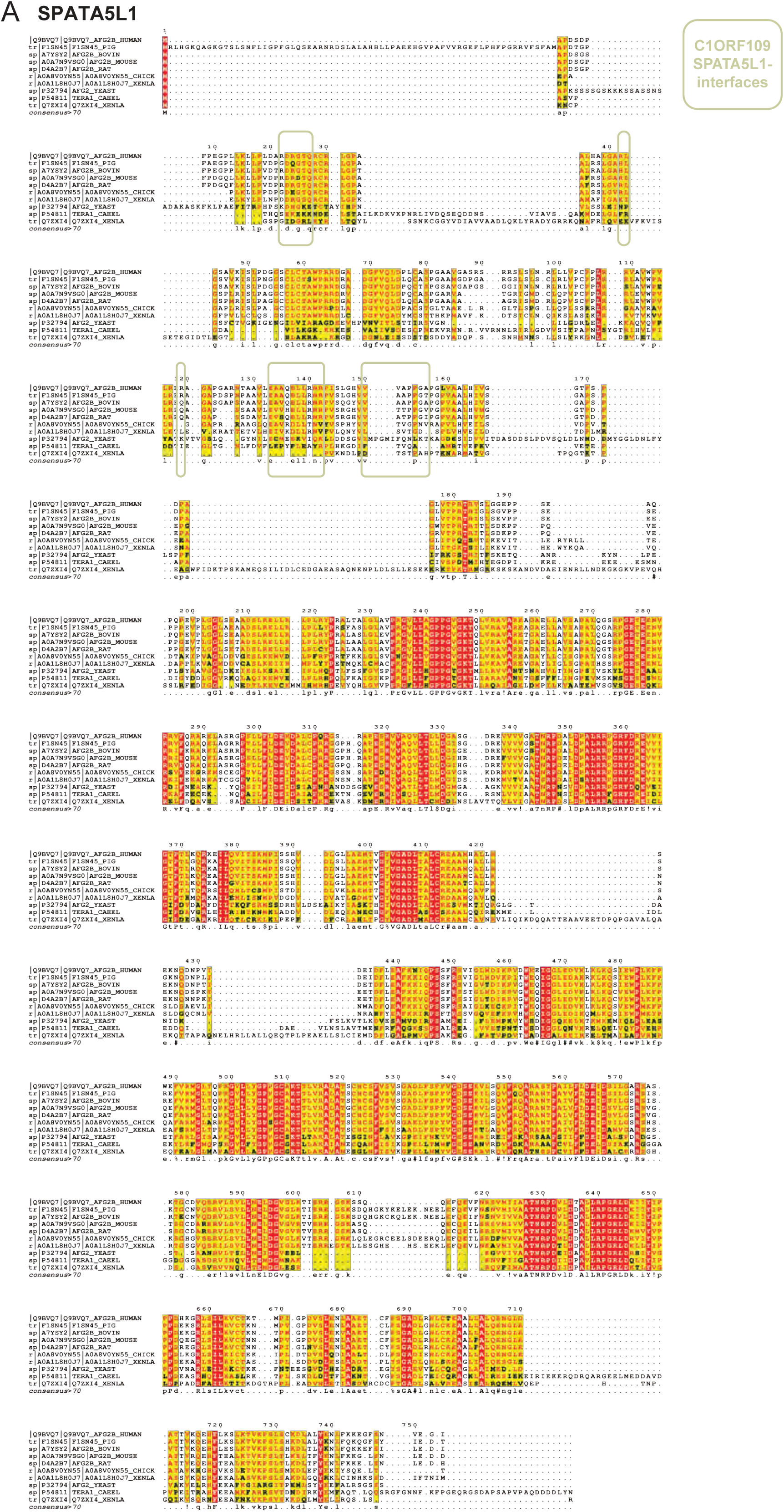
Multiple sequence alignment of SPATA5L1 wild-type protein sequences. Alignment created by T-coffee multiple sequence alignment server^63^ and ESPript v. 3.095^64^. Interface areas with cofactor CINP-C1ORF109, as determined by cryo-EM and structural modeling, are indicated.

**Figure S10.**
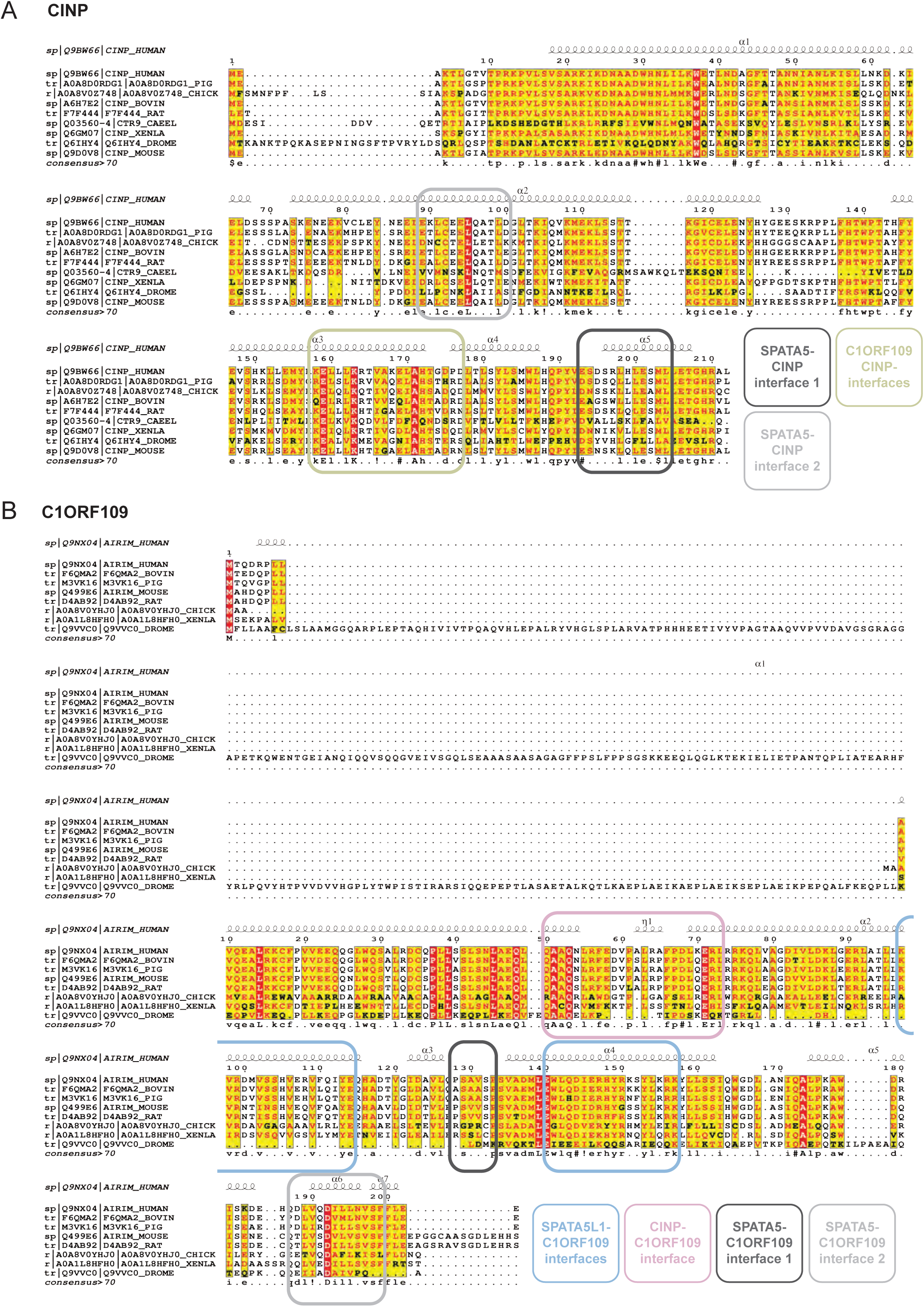
Multiple sequence alignment of CINP and C1ORF109 wild-type protein sequences. Alignment created by T-coffee multiple sequence alignment server^63^ and ESPript v. 3.095^64^. Interface areas with AAA+ ATPases SPATA5 and SPATA5L1, as determined by cryo-EM and structural modeling, are indicated.

**Figure S11.**
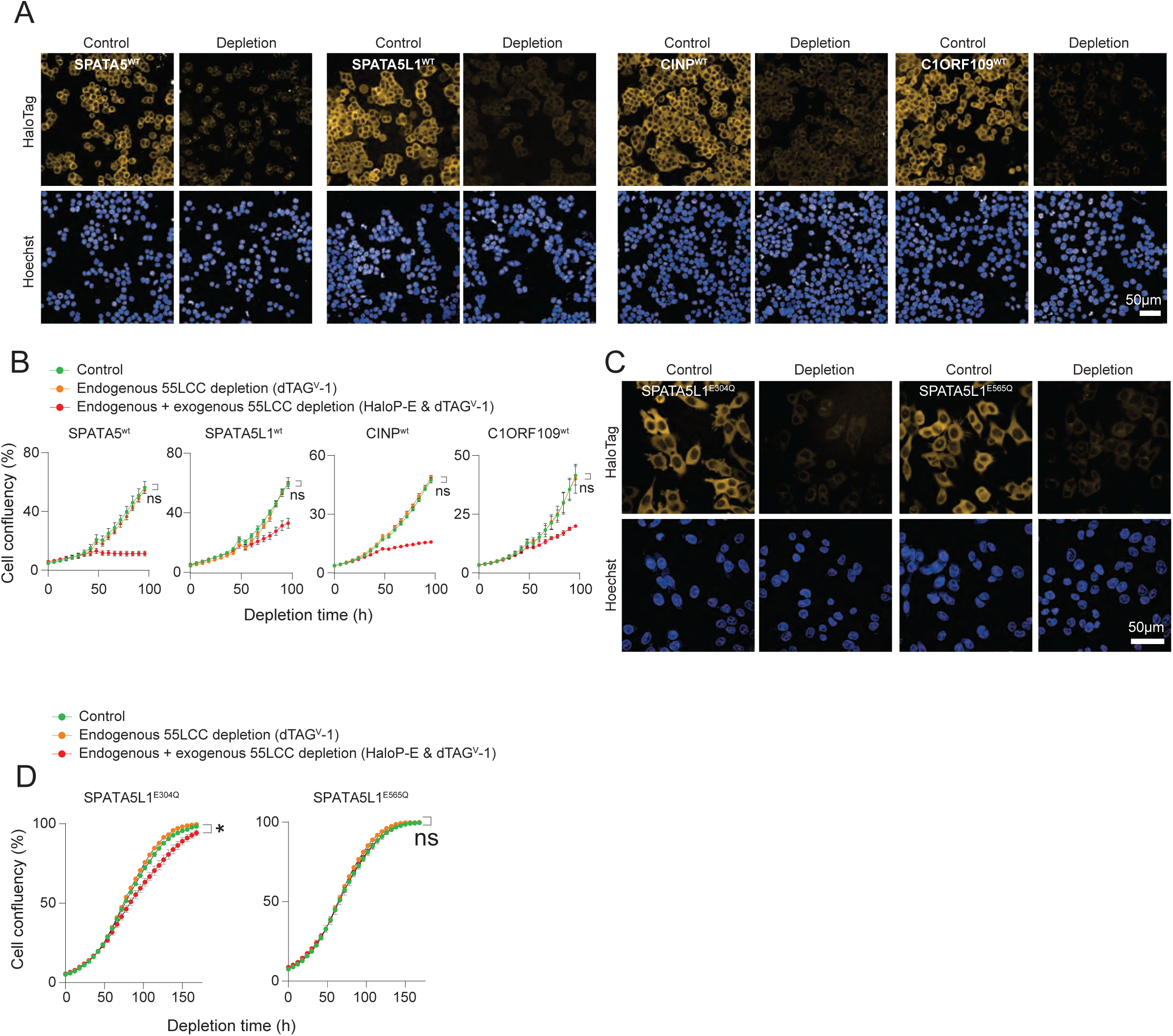
Functional validation of the strategy for acute depletion of endogenously expressed 55LCC constituents and rescue by exogenously expressed respective 55LCC constituents. (**A**) The cDNAs encoding the indicated 55LCC constituents were fused with HaloTag and the transgenes were inserted into the AAVS1 safe harbor locus of the corresponding dTAG-fused 55LCC constituents (i.e, HaloTag-fused SPATA5 ectopically expressed in cells expressing endogenous SPATA5^dTAG^). Micrographs showing HaloPROTAC-E-induced depletion (250 nM, 2 hours) of the indicated ectopically expressed 55LLC constituents. (**B**) Quantification of cell proliferation after the indicated treatments and cell lines. Cell proliferation analyzed by examining cell confluency by image-based analysis by Incucyte (*n*=3, Mean ± SEM). Statistical significance determined by 2-way ANOVA (Šidák-corrected). (**C**) Micrographs show expression and degradability of the indicated SPATA5L1 single ATPase-dead transgene variants following HaloPROTAC-E treatment (250 nM, 2 hours). (**D**) Cell proliferation analysis of the cells expressing the endogenous SPATA5L1^dTAG^ and the indicated SPATA5L1 single ATPase dead transgenes under the indicated treatment conditions and timepoints (*n*=3, Mean ± SEM). Cell proliferation and statistical significance quantified as specified in Figure 7B. (Statistical significance intervals indicated as: * p<0.05, ** p<0.01, *** p<0.001, **** p<0.0001.

## Supplementary Tables

**Supplementary Table 1** - Plasmids, gRNAs, oligos, cell lines

**Supplementary Table 2** - AP-MS data

**Supplementary Table 3** – DepMap gene essentiality scores

**Supplementary Table 4** – Curated list of proteins implicated in ribosome biogenesis

**Supplementary Table 5** - ΔCINP-C1ORF109 crystal structure data collection and refinement statistics.

**Supplementary Table 6** - MS DATA overview

**Supplementary Table 7** - MS settings

**Supplementary Table 8** – Protein interaction data unfiltered

**Supplementary Table 9** – Protein interaction network data filtered and curated

